# Correlative Synchrotron X-ray Microscopy Reveals Dose- and Division-Dependent Nanoparticle Redistribution in Macrophages

**DOI:** 10.64898/2026.02.21.707158

**Authors:** Isabella Scarpa, Renata S. Rabelo, Aline O. Pereira, Francine F. Fenandes, Flávia E. Galdino, Maiara F. Terra, Maria Harkiolaki, Florian Meneau, Carla C. Polo, André A. Thomaz, Ana J. Pérez Berná, Mateus B. Cardoso

## Abstract

Understanding the intracellular fate of nanoparticles is essential for designing safer and more effective nanomedicines, yet most studies rely on static observations and lack high-resolution, near-native volumetric information. Here, we establish a synchrotron-based correlative X-ray microscopy framework to investigate how fluorescent silica nanoparticles (SiNPs) redistribute within macrophages as a function of concentration and successive cell-division cycles. SiNPs were internalized by RAW 264.7 macrophages at different concentrations and analyzed using a synchrotron-based correlative X-ray microscopy workflow integrating cryogenic soft X-ray tomography (cryo-SXT), cryogenic structured illumination microscopy (cryo-SIM), and coherent X-ray ptychography, with confocal fluorescence microscopy used to establish population-level uptake tendencies. Cryo-SXT reveals a concentration-dependent redistribution of nanoparticle-containing vesicles from peripheral endosomes toward the perinuclear region, while correlative cryo-SIM confirms strict vesicular confinement, with no evidence of free nanoparticle diffusion into the nucleoplasm. At higher doses, nanoparticles approach the nuclear region via vesicles extending into nuclear-envelope invaginations, rather than by true nuclear entry. Successive cell divisions redistribute the intracellular nanoparticle load and promote stable perinuclear clustering, identifying a long-term sequestration route in macrophages. Coherent X-ray ptychography further reveals nanoscale deformations of the nuclear envelope associated with dense perinuclear vesicles. Together, these results establish synchrotron-based correlative X-ray microscopy as a mechanistic, multiscale platform for unveiling the dynamic intracellular fate of nanoparticles and providing mechanistic insight into their apparent nuclear localization.

## INTRODUCTION

The intracellular fate of engineered nanoparticles plays a central role in determining their safety, bioavailability, and therapeutic performance in nanomedicine ^1–3^. Their ability to cross biological barriers and interact with specific intracellular targets makes them powerful tools for therapeutic innovation^4,5^. However, the biological performance of nanoparticles depends not only on their composition and design but also on their intracellular fate - the dynamic sequence of events that determines how they traffic, transform, and interact with organelles after uptake^6–10^. Understanding this fate is critical for improving nanoparticle safety, bioavailability, and targeting efficiency. Among the wide variety of nanomaterials, silica nanoparticles (SiNPs) stand out as versatile platforms due to their tunable size, porosity, and surface functionalization, which enable fine control over drug loading and molecular interactions^2,11–13^. Numerous studies have shown that the adsorption of biomolecules, mostly proteins, onto the nanoparticle surface results in the formation of the protein corona^14,15^ plays a decisive role in defining the biological identity of nanomaterials, influencing their cellular uptake, distribution, cytotoxicity^16^ and overall biocompatibility^17,18^. Despite these advances, the complex and dynamic nature of nanoparticle–biological interactions remains poorly understood, limiting our ability to design nanoparticles with predictable and reproducible behavior across diverse biological environments^19^.

One of the main reasons for this limitation is the difficulty of accurately determining the precise intracellular localization and behavior of nanoparticles, as conventional imaging methods offer either insufficient spatial resolution or lack the necessary biological context. Fluorescence microscopy offers molecular specificity but relies on external labeling, thereby restricting access to intrinsic ultrastructural information^20,21^. Transmission electron microscopy (TEM) provides atomic-level resolution but requires ultrathin-sectioned samples, which prevents visualization of entire cells in their native architecture^22^. These constraints underscore the need for advanced nano-imaging approaches that combine volumetric, label-free, and high-resolution capabilities under near-physiological conditions. In this context, X-ray–based methods have gained attention for their ability to overcome many of the limitations of conventional microscopy by exploiting the strong penetration and intrinsic contrast of biological materials^23^. Although achieving high spatial resolution still demands precise optical design and fabrication, the advent of synchrotron light sources has opened new possibilities for extending both the resolution and sensitivity of X-ray imaging^24,25^.

These technological advances are implemented through distinct physical contrast mechanisms that define how synchrotron-based X-ray imaging interrogates cells, ranging from absorption to coherent diffraction. Particularly, cryogenic soft X-ray tomography (cryo-SXT)^26,27^ and X-ray ptychography^28^ show great promise for achieving high-resolution biological X-ray imaging. For example, cryo-SXT has enabled label-free mapping of cellular ultrastructure and nanoparticle localization in diverse systems, including gold nanoparticles^29,30^, iron oxide nanoparticles (SPIONs)^31^, and silica-based nanomaterials^32^, revealing their confinement in endocytic and lysosomal compartments and their progressive accumulation near the nucleus. Correlative approaches combining cryo-SXT with cryogenic fluorescence microscopy, as recently demonstrated in the literature^32^, have shown that molecular identity can be directly linked to three-dimensional ultrastructural context. ^33^In parallel, X-ray ptychography has demonstrated its ability to provide high-resolution, quantitative electron-density maps of biological specimens and to locate nanomaterials within cells with nanometer-scale sensitivity^33,34^. Despite these advances, most biological applications of synchrotron X-ray imaging remain primarily descriptive, focusing on short incubation times, single concentrations, and static snapshots of nanoparticle localization. Systematic investigations that exploit X-ray techniques to follow how nanoparticle distribution evolves as a function of dose, time, and cell-division dynamics are still largely missing, particularly in immune cells.

Here, we address this methodological and biological gap by establishing a synchrotron X-ray–based imaging framework to investigate how nanoparticle distribution evolves as a function of concentration and successive cell division cycles in macrophages. By integrating cryo-SXT and X-ray ptychography in 2D and combined with tomography within a single experimental strategy, we move beyond static localization and use X-ray imaging as a dynamic tool to monitor nanoparticle accumulation, vesicular trafficking, and long-term intracellular redistribution. This approach reveals a concentration-dependent pathway in which SiNPs larger than 100 nm can access the nuclear region at early time points and are subsequently redistributed through cell divisions. We then introduce a nuclear isolation protocol compatible with ptychographic imaging that enables targeted high-resolution analysis of nanoparticle–nucleus interactions. Beyond structural visualization, synchrotron X-ray techniques are established here as a central experimental platform for addressing time- and dose-dependent questions in nanomedicine, providing a methodological framework for studying nanoparticle dynamics within complex and highly phagocytic immune cells.

## RESULTS AND DISCUSSION

Fluorescent silica nanoparticles (SiNPs), doped with the dye ATTO 633, were successfully synthesized as spherical structures, with a mean diameter of 135 ± 16 nm determined from the analysis of 250 individual particles by STEM imaging ( Figure S1, Supporting Information). The obtained size range (<200 nm) is particularly relevant for biomedical applications, as it favors efficient cellular uptake and intracellular trafficking^35^. The overall experimental workflow is illustrated in Figure 1a, where SiNPs were coated with bovine serum albumin (BSA) to form a protein corona, thereby improving biocompatibility by reducing cytotoxicity while enhancing the cellular uptake^16,32^. Cells were then incubated with these BSA-coated SiNPs for controlled periods, washed, and analysed at different time points, enabling us to monitor the intracellular distribution of nanoparticles across successive cell divisions. The framework represents the experimental model adopted in this study, relating nanoparticle concentration to post-exposure time, during which increasing cell number reflects ongoing proliferation (Figure 1b). By systematically varying nanoparticle concentration and post-exposure time, we correlated the extent of SiNPs internalization with cell proliferation dynamics, providing mechanistic insight into the redistribution of nanoparticle load during the mitotic cycle.

**Figure 1.**
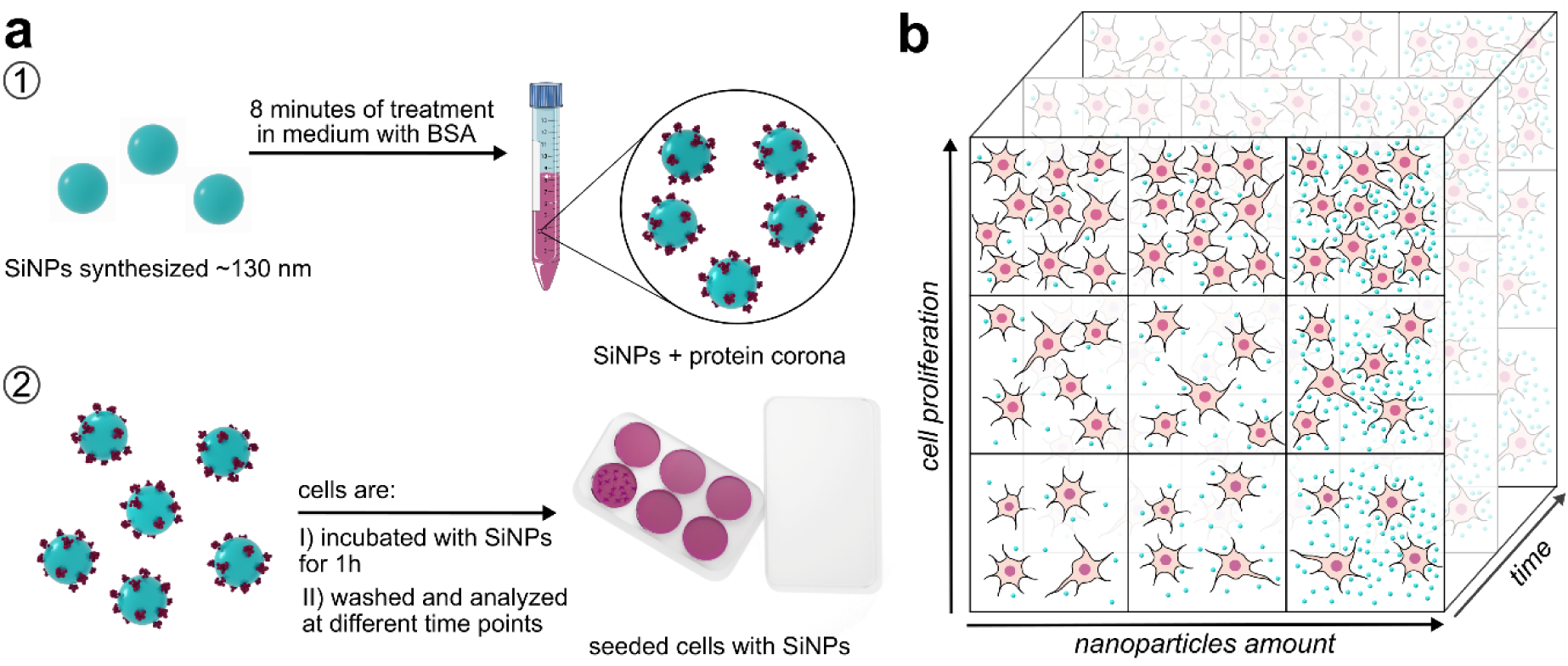
Experimental workflow and conceptual framework. a) 1- SiNPs are treated with BSA for 8 minutes to form a protein corona. 2 - SiNPs are incubated in RAW 264.7 cells for 1 h, subsequently washed with phosphate buffered saline (PBS), and analysed at different time points. b) Schematic representation of SiNPs distribution as a function of nanoparticle concentration and post-exposure time. The increase in cell number arises from proliferation along the time axis.

Understanding how nanoparticles interact with immune cells is essential for predicting their biological behavior and long-term *in vivo* fate. Regardless of their intended biomedical application, SiNPs inevitably encounter immune cells, which serve as the first line of recognition and clearance of foreign materials ^36,37^. To evaluate how nanoparticle load influences cellular responses, RAW 264.7 macrophages were exposed to three SiNPs concentrations (0.003, 0.03, and 0.3 mg/mL). This range was selected to capture potential dose-dependent effects on internalization efficiency, vesicular organization, and overall intracellular distribution ^38^. Flow cytometry (Figure S5 Supporting Information) was used to quantify nanoparticle uptake across the three tested concentrations. Cells were stained with propidium iodide (PI) while SiNPs (containing ATTO 633) were detected in the APC channel, enabling discrimination between nanoparticle-positive (ATTO+PI) and nanoparticle-free (PI) populations. At the lowest concentration (0.003 mg/mL), only 79.6% of cells were nanoparticle-positive (Figure S5c), indicating limited extracellular availability. This incomplete initial loading leads to progressive dilution of nanoparticles and a decrease in the fraction of nanoparticle-positive cells across successive divisions. In contrast, at 0.03 and 0.3 mg/mL, nearly all cells contained SiNPs at the initial time point (Figure S5a, b). This near-uniform loading ensures that most cells retain nanoparticles after division, maintaining a high fraction of nanoparticle-positive cells over multiple cell cycles. This strong concentration dependence reflects the high phagocytic activity of macrophages and their capacity to modulate uptake in response to particle abundance ^39,40^.

A multimodal imaging strategy was used to resolve how intracellular nanoparticle organization evolves with concentration. Confocal fluorescence microscopy provided an initial overview of nanoparticle distribution after 1 h of incubation (Figure 2a). The ATTO 633 signal from the particles increased progressively from 0.003 to 0.3 mg/mL, reflecting the higher nanoparticle availability in the extracellular medium and is consistent with the flow cytometry results. To solve the ultrastructural context of these fluorescent signals, cryo-SXT at B24 Beamline (Diamond) was employed under near-native conditions (Figure 2b). Tomographic slices revealed high-contrast dense objects, highlighted by orange arrows, corresponding to SiNPs encapsulated within vesicles, indicating that across all concentrations SiNPs were internalized via vesicular pathways^41,42^. However, their spatial organization strongly depended on nanoparticle dose. At low concentration, only a small vesicle containing SiNPs was detected in the cytoplasm. At intermediate concentration, multiple vesicles containing SiNPs were observed in the cytosol. Notably, at the highest concentration (0.3 mg/mL), large vesicular structures containing SiNPs were distributed throughout the cytosol and within the nuclear region. This concentration-dependent progression is summarized schematically in Figure 2c. Correlative cryogenic structured illumination microscopy (cryo-SIM) and cryo-SXT imaging further validated these observations (Figure 2d). Cryo-SIM identified SiNPs (red) within vesicular compartments that colocalized with lysosomal markers (blue), confirming their endolysosomal confinement^41^. Overlay with cryo-SXT showed that the colocalization of lysosomal and SiNP signals corresponds to the high-contrast structures observed in the tomograms. Also, both cryo-SXT and cryo-SIM revealed vesicle-encapsulated SiNPs located within the nuclear region. Cryo-SIM confirmed that these particles remained enclosed by vesicular membranes, excluding free diffusion into the nucleoplasm, as also confirmed by the fluorescence confocal (Figure 2a), where the white arrows point to the overlap of the nuclear and nanoparticle signals. While nuclear localization has previously been reported mainly for ultrasmall nanoparticles or systems engineered for nuclear targeting^43–45^, this observation demonstrates vesicle-encapsulated access of non-functionalized silica nanoparticles larger than 100 nm to the nuclear region, without nucleoplasmic entry. The combined X-ray and fluorescence data suggest that this phenomenon arises from vesicle-induced deformation and invagination of the nuclear envelope, rather than active transport through nuclear pores, consistent with mechanical stress–induced nuclear envelope remodeling^46^.

**Figure 2.**
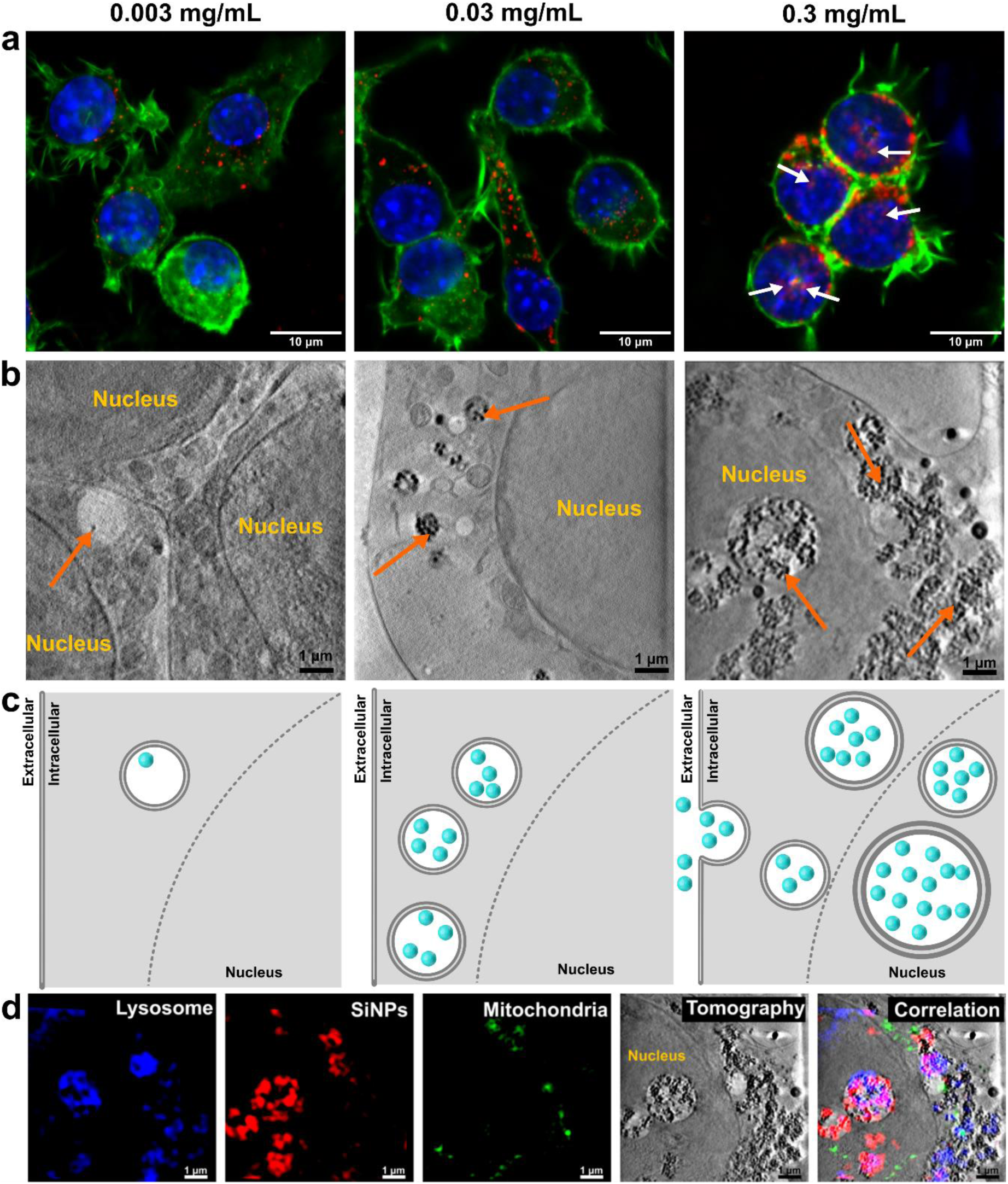
Internalization and intracellular distribution of SiNPs in RAW 264.7 cells at different concentrations. a) Fluorescence confocal micrographs of cells exposed to SiNPs at 0.003, 0.03, and 0.3 mg/mL (green: actin; blue: nucleus; red: SiNP). b) Slices from cryo-SXT tomographic reconstructions. Orange arrows indicate internalized SiNPs. c) Schematic representation illustrating the evolution of SiNP concentration inside macrophages as a function of exposure. d) Correlative imaging combining cryo-SIM and cryo-SXT shows lysosomes (blue), mitochondria (green), and SiNPs (red), followed by grayscale tomographic slices at 0.3 mg/mL and the merged image.

Building upon these concentration-dependent results, we next examined how cell proliferation dynamics influence nanoparticle redistribution across successive division cycles. Determining the doubling time (DT) was crucial to ensure experimental reproducibility, as cell growth rates can vary significantly between laboratories and culture conditions. Monitoring DT also allowed normalization of exposure periods to the cells’ biological rhythm, enabling comparison across equivalent proliferative stages. Fluorescence confocal and cryo-SXT imaging (Figure 3a–c) reveal a clear temporal evolution in SiNPs intracellular distribution at 0.3 mg/mL concentration over two division cycles. At the initial time (Figure 3a), SiNPs appear dispersed throughout the cytoplasm and within the nuclear region, as indicated by white arrows, with minimal clustering. After the first doubling (Figure 3b), fluorescence intensity increased and small perinuclear aggregates emerged, indicating progressive clustering of SiNPs, and cryo-SXT also revealed vesicles containing SiNPs relocating away from the nuclear region. By the second doubling (Figure 3c), SiNPs are predominantly concentrated around the nucleus, forming distinct high-intensity clusters consistent with late endosomal/lysosomal trafficking^9,41,42^. Segmented cryo-SXT volumes provide complementary ultrastructural confirmation of these patterns, highlighting both the redistribution of SiNP-containing vesicles during cell division and their preferential perinuclear accumulation. Correlative cryo-SIM and cryo-SXT analyses (Figure 3d) further elucidate the subcellular context, revealing the spatial relationship between SiNPs (red) and lysosomes (blue). After two division cycles, SiNPs remain confined within vesicular membranes in the perinuclear region^47^ suggesting that the observed redistribution results from active vesicular transport and maturation rather than passive diffusion.

**Figure 3.**
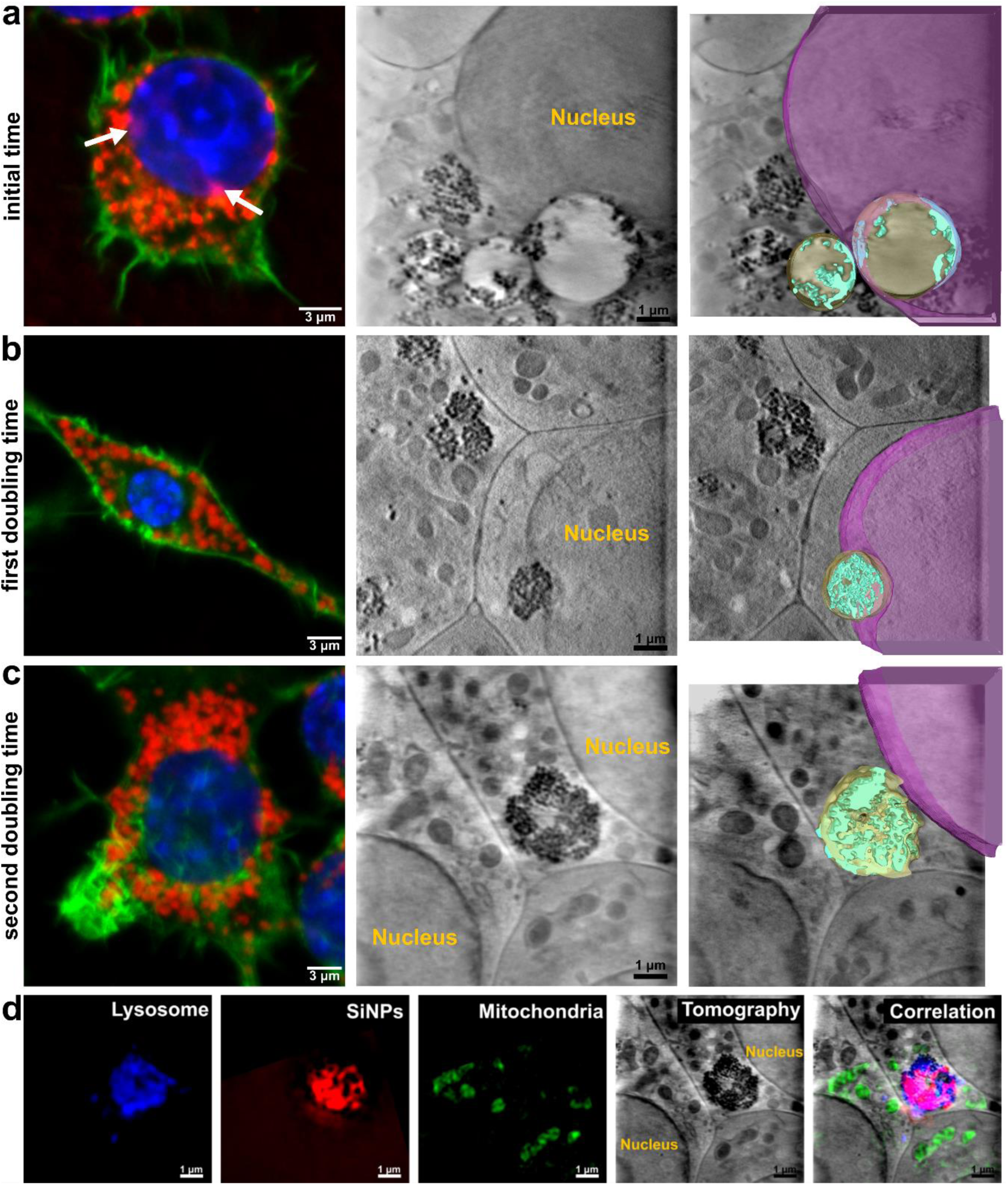
Multimodal imaging of intracellular SiNP distribution across two cell-division cycles in macrophages.(a–c) Fluorescence confocal micrographs (actin, green; nucleus, blue; SiNPs, red), corresponding cryo–SXT tomographic slices, and segmented volumes corresponding to the evolution of intracellular SiNP localization over time: a) initial time, b) first doubling time, and c) second doubling time. d) Correlative cryo–SIM and cryo–SXT imaging of lysosomes (blue), SiNPs (red), and mitochondria (green), followed by grayscale tomographic slices at the second doubling time and the merged correlation image.

Furthermore, X-ray mosaic projections of whole cells obtained at Mistral Beamline (Alba Synchrotron) (Figure 4a–c) extend this analysis to near–whole-cell volumes, showing that at the initial time SiNPs-containing vesicles are broadly distributed throughout the cytoplasm, whereas over successive division cycles they progressively concentrate toward the perinuclear region. Three-dimensional rendered and segmented volumes obtained from stitched cryo-SXT tomograms (Figure 4d–f) confirm this redistribution at the ultrastructural level, revealing a gradual relocation of vesicle-confined SiNPs toward the nuclear periphery. The schematic representations (Figure 4g–i) summarize this dynamic process, highlighting the progressive perinuclear aggregation across division stages. Together, this whole-cell perspective rules out local sampling effects and confirms that the observed perinuclear accumulation reflects a global intracellular reorganization rather than a region-specific phenomenon. This localization likely reflects a combination of endocytic processing, vesicle trafficking, and sequestration mechanisms that preserve cellular homeostasis during proliferation^47^. Together, these findings demonstrate that SiNP intracellular organization is dynamic and responsive to cell division, with progressive perinuclear accumulation reflecting coordinated endocytic activity and stable vesicular entrapment^38^. Such behavior provides structural insight into how proliferative cells adapt to nanoparticle exposure, revealing vesicular confinement as a key mechanism governing long-term nanoparticle retention and intracellular fate.

**Figure 4.**
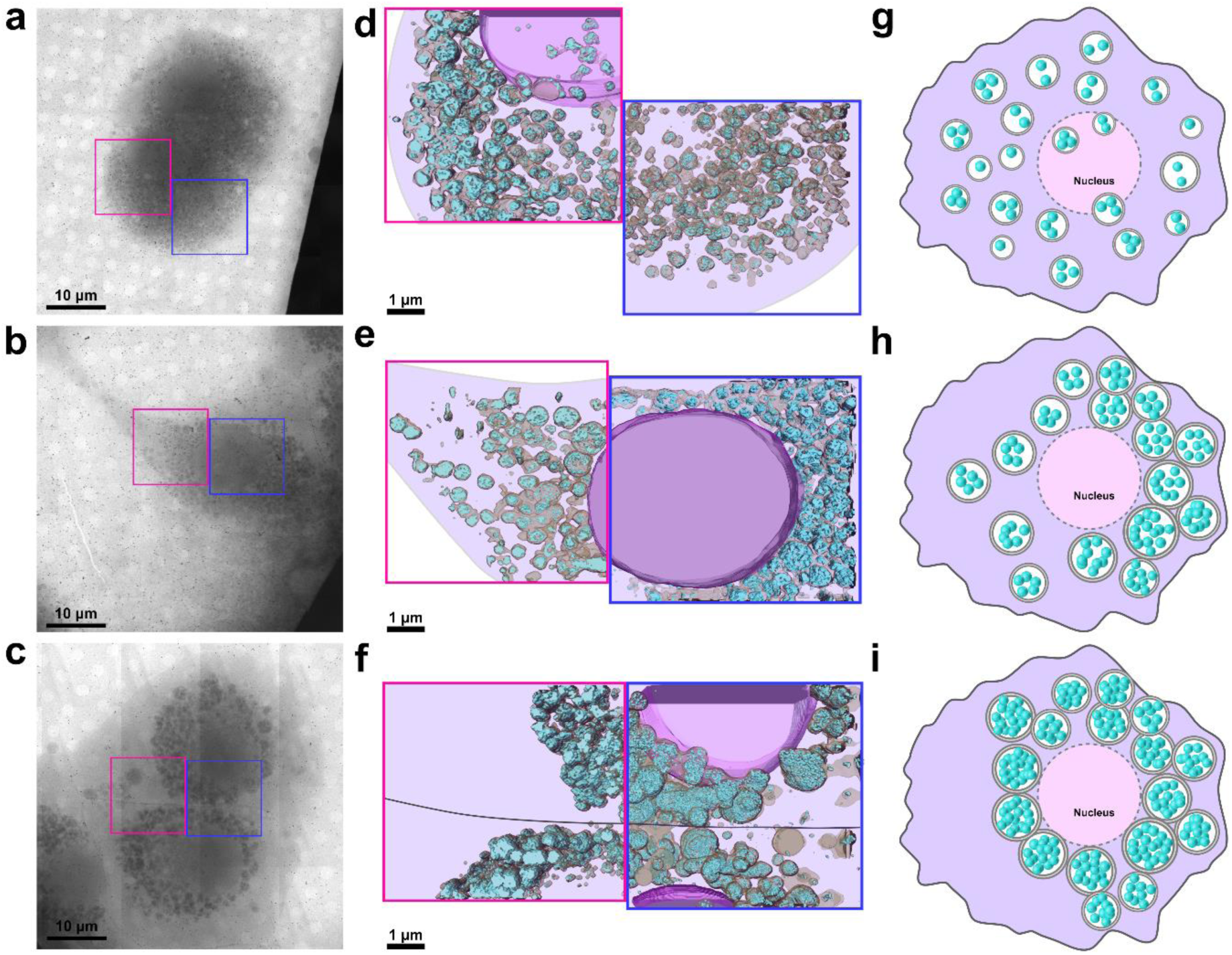
Time-dependent intracellular distribution of SiNPs at 0.3 mg/mL. (a–c) X-ray mosaic projections of whole cells at a) initial time, b) first doubling time, and c) second doubling time. Pink and blue squares indicate adjacent, partially overlapping regions used for tomographic reconstruction. (d–f) Three-dimensional rendered and segmented volumes obtained from stitched tomograms corresponding to the regions highlighted in panels (a–c) at d) initial time, e) first doubling time, and f) second doubling time. Highlighted structures include the cell nucleus (pink), vesicles (gold), and SiNPs (cyan); the cell boundary is indicated by a purple contour.(g–i) Schematic representations summarizing the progressive perinuclear aggregation of SiNPs-containing vesicles at g) initial time, h) first doubling time, and i) second doubling time.

We next employed X-ray ptychography, an advanced configuration of coherent diffractive imaging (CDI), to investigate these interactions at nanoscale resolution, particularly under higher SiNP concentrations. This technique provides quantitative, label-free imaging based on phase shifts induced by variations in the specimen’s electron density, enabling visualization of dense intracellular regions that are otherwise inaccessible to conventional microscopy. To enable this analysis, a nuclear isolation protocol^48,49^ was developed to lyse the plasma membrane while preserving the nuclear envelope selectively. It provides an essential means to investigate the potential structural influence of nanoparticle exposure on the nucleus. The optimized treatment with 1% Triton X-100 for 3 min, followed by glutaraldehyde fixation and critical-point drying, provided optimal preservation of nuclear integrity and compatibility with coherent X-ray diffraction imaging (Figure 5a). The experimental optical layout of the ptychography setup at Cateretê Beamline (Sirius) is schematically illustrated in Figure 5b. Unlike conventional lens-based imaging, X-ray ptychography reconstructs the complex electron density of a sample from overlapping diffraction patterns acquired during a scanning sequence^28^. Each illuminated position generates an interference pattern (“speckle”), which is computationally processed through iterative phase-retrieval algorithms^50^ such as the Ptychographic Iterative Engine (PIE)^51^ or the Difference Map (DM)^52^ to recover both amplitude and phase information. When extended to ptychographic X-ray computed tomography (PXCT)^53^, this approach enables three-dimensional visualization of entire cells with spatial resolutions reaching tens of nanometers. Despite its high resolving power, the application of X-ray ptychography to biological systems remains challenging due to the intrinsically low scattering contrast of biological materials and their susceptibility to radiation damage. However, recent advances in beam coherence and scanning stability (enabled by fourth-generation synchrotron light sources such as SIRIUS)^24,54,55^ have begun to overcome these limitations, allowing high-resolution imaging of intact cells. In this context, our study extends the use of ptychography to probe nanoparticle–nucleus interactions in immune cells, providing a three-dimensional, nanoscale view of how intracellular SiNPs influence nuclear architecture.

**Figure 5.**
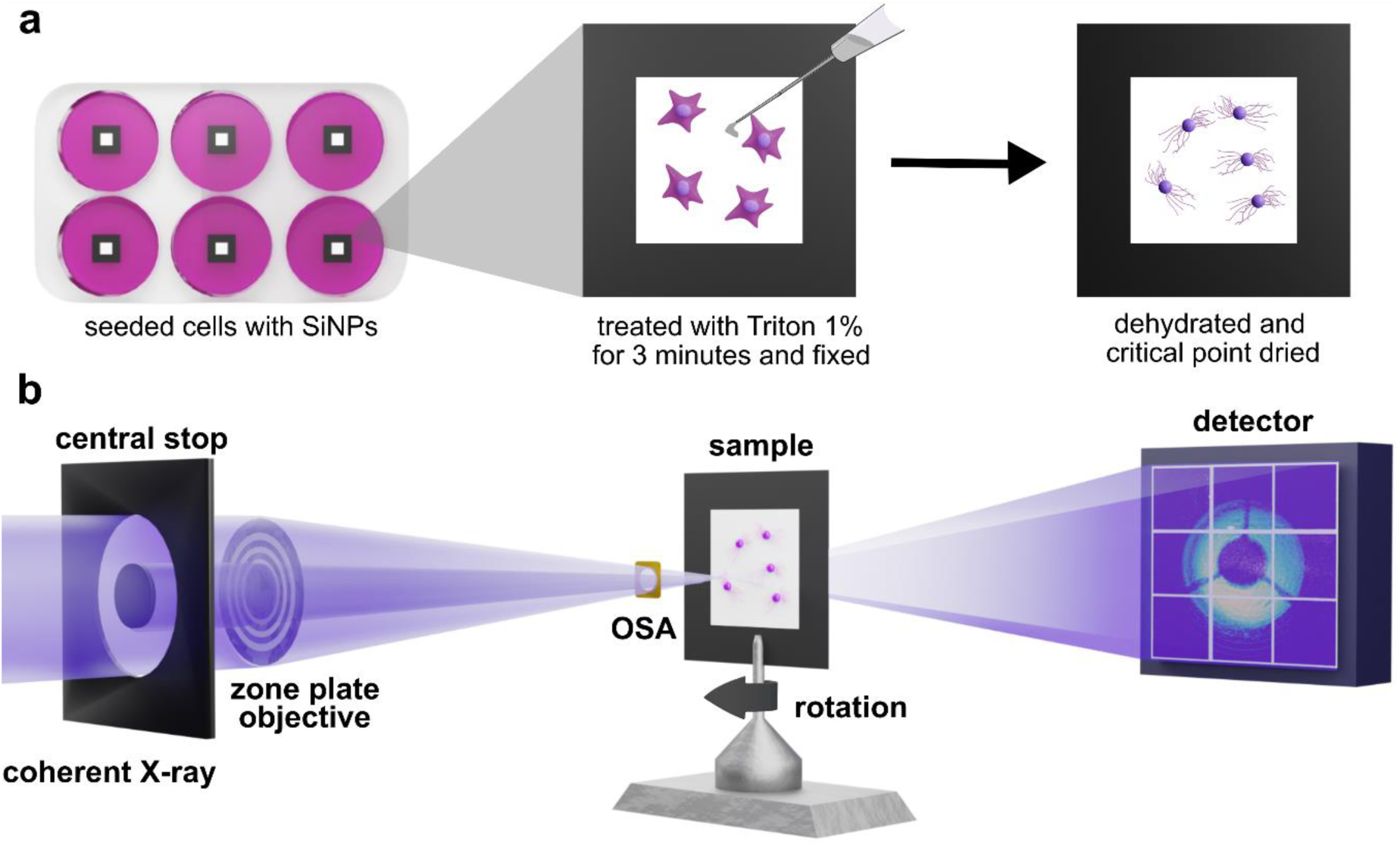
Experimental workflow and ptychographic imaging setup. a) Stepwise preparation of SiNP-treated cells for ptychography, including decellularization with 1% Triton for 3 min, fixation, dehydration, and critical point drying. b) Schematic representation of the ptychography optical layout, showing the path of the coherent X-ray beam through the sample membrane toward the detector.

Despite the higher radiation dose required, this approach enables detailed visualization of nuclear morphology, subnuclear organization, and intracellular nanoparticle distribution at different exposure times (Figure 6). The time-dependent intracellular distribution of SiNPs revealed by X-ray ptychography is summarized in schematic representations of the spatial organization shown in (Figure 6a, e, i, m). Two-dimensional ptychographic reconstructions revealed clear contrasts between the nucleus and cytoplasm (Figure 6b, f, j, n), with SiNPs appearing (Figure 6f, j, n) as high-density domains surrounding and occasionally deforming the nuclear envelope. These findings are consistent with the previously observed vesicle-mediated internalization pathways, suggesting that mechanical interactions between enlarged endosomal vesicles and the nucleus may drive local invaginations of the nuclear membrane (Figure 6e, f, g, h). Three-dimensional ptychographic tomograms (Figure 6h, l, p) further confirmed the progressive perinuclear accumulation of SiNPs over time^47^. After 1 h of incubation (Figure 6e, f, g, h), nanoparticles were observed near the nuclear periphery and within the nuclear region (MovieS6, Supporting Information), while after 9 h (Figure 6i, j, k, l; MovieS7) and 18 h (Figures 6m, n, o, p; MovieS8), dense clusters formed around the nucleus, in agreement with the confocal and cryo-SXT data. These results indicate that nanoparticle redistribution during successive cell-division cycles is accompanied by structural remodeling of the perinuclear compartment, reinforcing the notion of persistent vesicular entrapment rather than direct nuclear diffusion. Such PXCT analyses underscore the potential of coherent X-ray imaging to elucidate nanoscale interactions at the bio-nano interface with unprecedented clarity.

**Figure 6.**
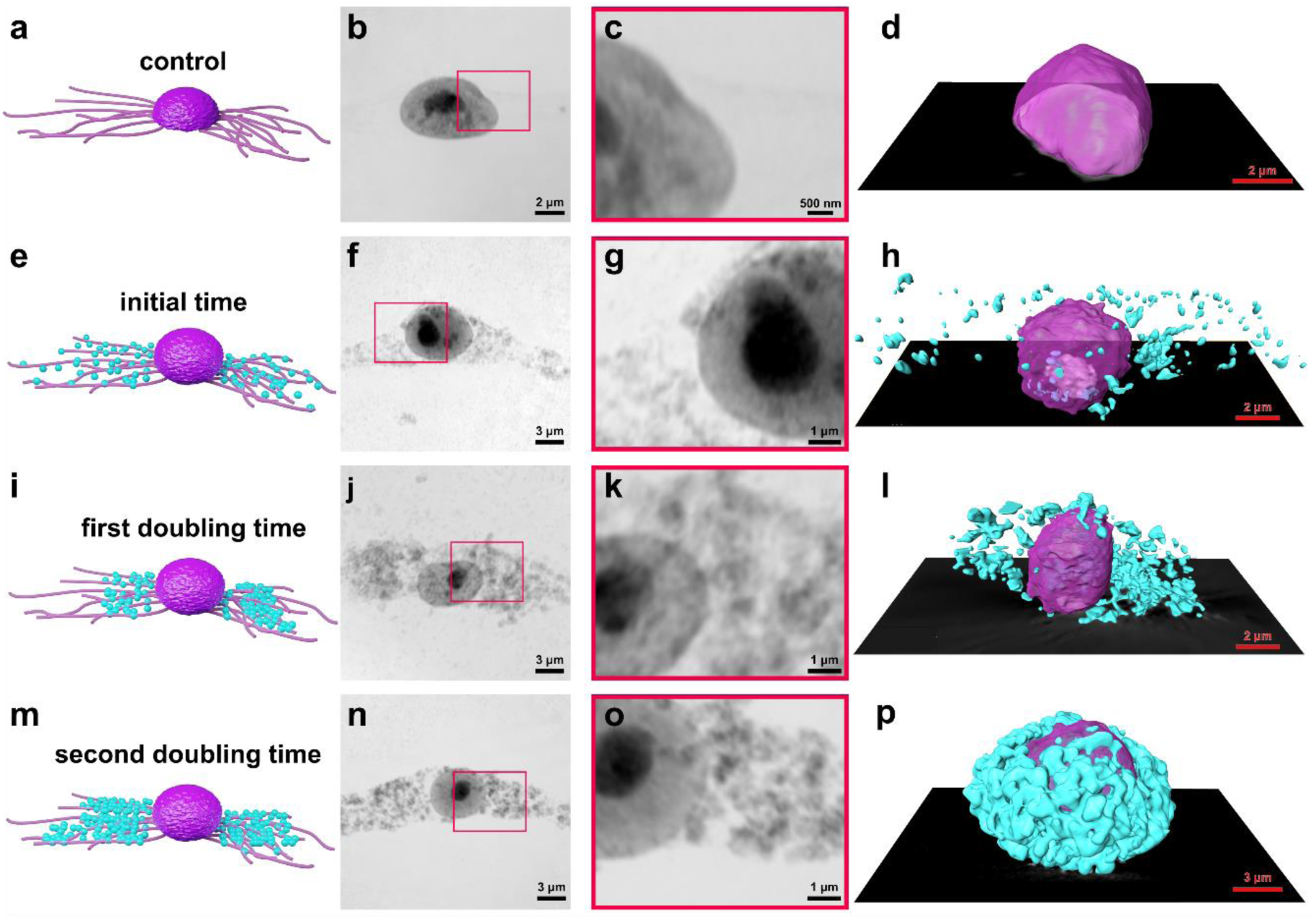
Time-dependent intracellular distribution of SiNPs revealed by X-ray ptychography. Schematic representation of the spatial distribution of silica nanoparticles (cyan) around the nucleus (purple) at each time point: a) Control cell; e) 0.3 mg/mL after 1 h; i) 0.3 mg/mL after 9 h (first doubling time); m) 0.3 mg/mL after 18 h (second doubling time). Two-dimensional ptychographic reconstructions of nucleus of RAW 264.7 cells: : b) Control cell; f) 0.3 mg/mL after 1 h; j) 0.3 mg/mL after 9 h; n) 0.3 mg/mL after 18 h. Magnified views of the boxed regions in two-dimensional ptychographic reconstructions, highlighting nuclear morphology and nanoparticle localization: c) Control cell; g) 0.3 mg/mL after 1 h; k) 0.3 mg/mL after 9 h; o) 0.3 mg/mL after 18 h. Three-dimensional renderings of the tomographic reconstructions: d) Control cell; h) 0.3 mg/mL after 1 h; l) 0.3 mg/mL after 9 h; p) 0.3 mg/mL after 18 h.

## CONCLUSIONS

In this work, we establish synchrotron-based correlative X-ray microscopy as a central platform to unravel the dynamic intracellular fate of nanoparticles in immune cells. By combining correlative cryo-soft X-ray tomography with X-ray ptychography, we show that the intracellular distribution of silica nanoparticles is not static but undergoes a dose - and division-dependent redistribution in macrophages. Cryo-SXT reveals volumetric, label-free maps of nanoparticle-containing vesicles, demonstrating concentration-dependent access to the nuclear region and progressive perinuclear accumulation over successive divisions. Correlative cryo-SIM confirms the endolysosomal nature of these compartments and excludes free diffusion into the nucleoplasm, while large-volume reconstructions show that this redistribution reflects a global intracellular reorganization. 2D ptychography and PXCT extends these observations to the nanoscale, revealing vesicle-induced deformation of the nuclear envelope. Together, these results identify vesicular confinement, mitotic redistribution, and perinuclear accumulation as key mechanisms governing long-term nanoparticle retention in macrophages. More broadly, this work positions synchrotron-based X-ray imaging as a dynamic analytical platform for investigating time- and dose-dependent nano–bio interactions across complex cellular systems.

## MATERIALS AND METHODS

### Synthesis and purification of fluorescent SiNPs

The nanoparticles were synthesized using a modified Stöber method^56^. One milligram of ATTO633-NHS-ester (λex/em: 630/651 nm) was dissolved in 800 µL of anhydrous dimethylformamide (DMF). Four hundred µL of this solution were added to 2 mL of anhydrous ethanol, followed by 9.6 µL of (3-aminopropyl) triethoxysilane (APTES). The reaction was maintained at room temperature with constant stirring for 20 hours, resulting in ATTO633-APTES. Then, 7 mL of ammonium hydroxide solution was added to 120 mL of ethanol and kept under stirring at room temperature. After 30 min, 2.4096 mL of ATTO633-APTES and 2.5 mL of tetraethoxysilane (TEOS) were added. After 3 hours, an additional 2.5 mL aliquot of TEOS was added, and the reaction was maintained under stirring for 24 hours. The nanoparticles were purified by centrifugation. The final volume of the synthesis was divided into three Falcon tubes (approximately 40 m L each) and centrifuged at 10,000 rpm for 15 min at 20 °C. After centrifugation, the supernatant was discarded, and 30 mL of ethanol was added to each Falcon tube. The nanoparticles were resuspended using a vortex and sonicated for 30 min. After another centrifugation step, the supernatant was discarded, and the nanoparticles were resuspended in 40 mL of ultrapure water. They were then placed in an ultrasonic bath for 30 min. This washing procedure with water was repeated four times. The final suspension was stored at 8°C, and the concentration was determined by gravimetry.

### Confocal fluorescence microscopy of RAW 264.7

RAW 264.7 macrophages cells were seeded at a density of 1.0 × 10^5^ cells per well in 6-well plates containing sterile glass coverslips. Prior to cell seeding, coverslips were sequentially cleaned under agitation in nitric acid (HNO₃, 15 min), rinsed thoroughly with distilled water, incubated in 1 N NaOH (15 min), rinsed again with distilled water, and treated with 70% ethanol (30 min). Coverslips were air-dried on filter paper, transferred to glass containers, and sterilized by autoclaving. SiNPs were used at concentrations of 0.3, 0.03, and 0.003 mg/mL. Prior to cell exposure, nanoparticles were incubated for 8 min in DMEM containing 1% BSA. Cells after 24 h were treated with the nanoparticle suspensions for 1 h at 37 °C. After incubation, cells, including untreated controls, were washed with PBS and fixed with 250 µL of 4% paraformaldehyde per coverslip for 10 min at room temperature. Fixation was followed by three washes with cold PBS (5 min each). This procedure was repeated for experimental time points of 9 h (calculated DT for BR-cultured cells) and 18 h following nanoparticle incubation at a concentration of 0.3 mg/mL. For permeabilization and blocking, coverslips were incubated with 250 µL of a solution containing Triton X-100 and BSA for 60 min at room temperature, followed by three additional washes with cold PBS (5 min each). The actin cytoskeleton and cell nuclei were subsequently labeled with Alexa Fluor 488–conjugated phalloidin and DAPI, respectively. Fluorescence imaging was performed using an inverted Zeiss LSM confocal microscope equipped with an Airyscan detector, located at INFABiC (National Institute of Science and Technology in Applied Photonics to Cell Biology), University of Campinas (UNICAMP). Images were acquired using a 63× oil immersion objective (NA 1.4).

### Cryo-SXT and Cryo-SIM of RAW 264.7 at Diamond Light Source

Gold and carbon grids (Quantifoil AU G200F1 finder) were cleaned with 70% ethanol, rinsed with PBS, and placed in 6-well plates containing 10% FBS in water for 24 h, with the carbon side facing upward. RAW 264.7 cells (250,000 cells/well) were seeded onto the carbon side of the grids for 24 h (37 °C, 5% CO₂). SiNPs at concentrations of 0.3, 0.03, and 0.003 mg/mL were preincubated with 1% BSA in DMEM for 8 min. Cells were then treated with the nanoparticles for 1 h and washed with PBS. Control samples were prepared without nanoparticle treatment. Accordingly, after 1 h, 16 h (calculated DT for UK-cultured cells), and 32 h, cells were treated for 30 min with LysoTracker Blue 100 nM (λex/em 373/422 nm) and MitoTracker Green 100 nM (λex/em 490/516 nm), diluted in DMEM + 10% FBS. The cells were then vitrified immediately in liquid ethane, without prior washing, and kept in liquid nitrogen until the analyses were performed. All protocols pertaining to this sample preparation have been described elsewhere^57^. Firstly, the grids were mapped using the Zeiss Axio Imager M2 coupled to a Linkam cryostage. Subsequently, images were collected using the cryo-SIM, at beamline B24 at the Diamond Synchrotron Light Source (Didcot, UK) to determine the location of lysosomes and nanoparticles. The samples were kept on a cryo-stage (CMS196M, Linkam Scientific), and a 100x magnification objective, numerical aperture 0.9, and 2 mm working distance were used (Nikon). Subsequently, analyses by cryo-SXT of these same regions were performed on an UltraXRM-S220C microscope (Carl Zeiss X-ray Microscopy Inc.), also on beamline B24 at the Diamond Synchrotron Light Source (Didcot, UK). This analysis was performed in the energy range known as the water window (284 eV - 543 eV) at 520 eV. A depth of focus of 1 µm was selected, achieving a resolution of 25 nm. The reconstructed tomograms were segmented using Avizo software (Thermo Fisher Scientific). This software was employed for surface rendering and three-dimensional visualization of the intracellular distribution of nanoparticles. Segmentation was performed manually based on the reconstructed tomograms.

### Cryo-SXT of RAW 264.7 at Alba Synchrotron

Gold–carbon grids were cleaned by glow discharge and exposed to UV radiation for 30 min. Subsequently, the grids were placed carbon-side up in 60 mm culture dishes containing FBS diluted in water and incubated for 1 h. RAW 264.7 cells (300,000 cells per well) were seeded onto the carbon side of the grids and cultured for 24 h at 37 °C under 5% CO₂. SiNPs at a concentration of 0.3 mg/mL were pre-incubated with 1% BSA in DMEM for 8 min. Cells were then treated with the nanoparticles for 1 h and washed with PBS. Control samples were prepared under the same conditions without nanoparticle treatment. The DT of RAW 264.7 cells cultured in Barcelona was previously determined to be 15 h. Based on this value, samples were collected after 1 h, 15 h, and 30 h, in addition to untreated controls. At each time point, samples were washed with PBS to remove excess medium and ensure optimal X-ray transmission. For tomographic alignment, 1 µL of a 100 nm gold nanoparticle suspension (3.60 × 10^8^ particles/mL, EMGC100, BBI Group, Cardiff, UK) was added onto the grids. The grids were rapidly vitrified by plunge freezing in liquid ethane using a Leica EMCPC system. Frozen grids were imaged and selected using a LINKAM CMS196 stage mounted on a Zeiss Axio Scope fluorescence microscope. The vitrified grids were subsequently stored in liquid nitrogen until imaging. Cryo-SXT was performed at the Mistral beamline of the ALBA Synchrotron using an UltraXRM-S220C microscope (Carl Zeiss X-ray Microscopy Inc.)^58,59^. Tomographic data were collected at 520 eV, with exposure times of 1–2 s per projection. Projection images acquired at different sample orientations were computationally combined to generate three - dimensional (3D) reconstructions of whole-cell subcellular ultrastructure^60^. A tilt series was acquired for each cell area using an angular step of 1° over a ±70° range, employing a Fresnel zone plate (FZP) with a 40 nm outermost zone width and an effective pixel size of 13 nm. Each transmission projection image of the tilt series was normalized using flat-field images, accounting for exposure time and storage ring current. Wiener deconvolution, considering the experimental impulse response of the optical system, was applied to the normalized data to enhance image quality^61^. The Napierian logarithm was then used to reconstruct the linear absorption coefficient (LAC). The resulting image stacks were loaded into IMOD software^62^, and individual projections were aligned to a common tilt axis using the 100 nm gold nanoparticles as fiducial markers. Aligned stacks were subsequently reconstructed using algebraic reconstruction techniques (ART)^63^. Tomographic reconstructions were segmented manually using Avizo software (Thermo Fisher Scientific) based on the reconstructed tomograms.

### 2D Ptychography and PXCT

RAW 264.7 cells were cultured in DMEM supplemented with 10% FBS at 37 °C in a 5% CO₂ atmosphere. Cells were seeded at a density of 10,000 cells per well in 6-well plates onto 100 nm-thick silicon nitride membranes, previously sterilized. After 24hrs they were then incubated with SiNPs coated with 1% BSA at a concentration of 0.3 mg/mL for 1 h. After this internalization period, cells were washed with PBS and maintained in DMEM. Untreated cells were used as controls. After 9 h and 18 h, the cells were washed with PBS and treated with 1% Triton for 3 min to lyse the plasma membrane while preserving intact nuclei. Cells were subsequently fixed with 2.5% glutaraldehyde in 0.1 M cacodylate buffer for 24 h, washed with 0.1 M cacodylate buffer and progressively dehydrated in a graded ethanol series (15%, 30%, 50%, 80%, 90%, 100%), followed by critical point drying with CO₂. Samples were first screened under an optical microscope to identify the most suitable regions for three-dimensional imaging, which was performed at the Cateretê beamline of the LNLS (Brazilian Synchrotron Light Laboratory). The area to be imaged was selected using the Arinax on-axis optical microscope. The 2D ptychographic and PXCT were carried out at the energy of 6 keV. The sample was illuminated by a beam defined by a set of central stop (CS), Fresnel Zone Plate (FZP) (50 um diameter and outermost zone width of 50 nm) and an order sorting aperture (OSA). The sample was placed 2.4 mm from the FZP focus, resulting in a beam size of 10 um, which scanned the sample with steps of 1.75 µm. The acquisition of each scanning point was 25 ms. The scattered X-rays were detected the in vacuum PiMega 540d detector (PiTec-LNLS, 55 um pixel size) positioned 7 m downstream from the sample. Bidimensional projections and tomographic reconstructions were performed using the PtychoShelves package^64^. Each projection was reconstructed into two-dimensional (2D) images using 500 iterations of the difference map (DM) algorithm. Subsequently, the set of 2D reconstruction were assembled to reconstruct the tomographic image, using Filter Back Projection (FBP) algorithm. Tomographic reconstructions were segmented using Avizo software (Thermo Fisher Scientific).

## Supporting information

SupportingInformation_001

## ACKNOWLEDGMENTS

The authors gratefully acknowledge the financial support provided by the Fundação de Amparo à Pesquisa do Estado de São Paulo (FAPESP – Processes 2021/12071-6, 2023/00103-6, 2024/00989-7). The authors also thank the Electron Microscopy Laboratory of LNNano for access to the electron microscopy facilities (Proposal SEM-20233386), LNBio for access to the flow cytometer (Proposal 20231954). We thank the access to equipment and assistance provided by the National Institute of Science and Technology on Photonics Applied to Cell Biology (INFABIC) at the State University of Campinas; INFABIC is co-funded by Fundação de Amparo a Pesquisa do Estado de São Paulo (FAPESP) (2014/50938-8) and Conselho Nacional de Desenvolvimento Cientifico e Tecnológico (CNPq) (465699/2014-6) for the access to the confocal fluorescence microscopy, LNLS for access to ptychography at the Caterete beamline (20250854 and 20250988), Diamond Light Source for access to cryo-SIM and cryo-SXT at the B24 beamline (BI34928), and Alba Synchrotron for access to cryo-SXT at the Mistral beamline (2024078502). The authors further acknowledge iNext (PID 24410 and VID 43520) for financial support during the experiments performed at Diamond Light Source. Special thanks are due to Vitor B. Pelgati for his assistance with confocal fluorescence microscopy, to Jessica do N. Faria for support with sample preparation for cryo-SXT experiments at the ALBA Synchrotron and to Archana Jadhav and Kamal L. Nahas for experimental support at Diamond Light Source. The authors also appreciate the contributions of Tiago A. Kalile and Pedro H. Z. Guidolim to the ptychography experiments.

