## SupportingInformation_001 for "Correlative Synchrotron X-ray Microscopy Reveals Dose- and Division-Dependent Nanoparticle Redistribution in Macrophages"

### Materials and Methods

#### *Characterization of nanoparticles*

The shape and size distribution of the nanoparticles were evaluated using STEM and dynamic light scattering (DLS). STEM images of SiNPs were obtained using SEM-FEI Inspect F50 (30 kV) microscope, at the Brazilian Nanotechnology National Laboratory (LNNano - CNPEM, Brazil). Samples were prepared by applying 5 µL of nanoparticle suspension to a copper microscopy grid and removing excess sample with filter paper after 10 minutes. Over 250 nanoparticles were analyzed for size distribution, and measurements were performed manually using ImageJ software. The hydrodynamic diameter ($D_H$) and polydispersity of 1 $g.L^{-1}$ SiNP@ATTO in water were measured using a Malvern Zetasizer ZS (Malvern Instruments Ltd., UK) at a 173° scattering angle and a wavelength of 633 nm.

#### *Cell doubling time*

To analyze the effect of cell doubling time, the doubling time (DT) was calculated using the following formula (Animal Cell Culture Guide):

$$DT = \frac{Tln2}{\ln\left(\frac{xt}{x0}\right)}$$

where *T* is the elapsed time (h), $x_0$ the initial cell concentration at plating, and $x_t$ the final concentration at time *T*.

***Fluorescence Spectroscopy***

The fluorescence of nanoparticles containing ATTO 633 was verified using fluorescence spectroscopy. To confirm the entrapment of ATTO 633 molecules within the SiNPs, a 100μL of fluorescent SiNPs at a concentration of 1 mg/mL was subjected to centrifugation for 15 minutes at 14.000 rpm. The fluorescence of both the supernatant and the precipitated nanoparticles, which were subsequently redispersed in 100 μL of water, was excited at 600 nm, and emission in the range of 618 to 840 nm was detected. Fluorescence measurements were performed using a Varioskan™ LUX multimode microplate reader (Thermo Fisher Scientific, model 3020-82689).

***Protein Corona***

To evaluate protein corona formation, the $D_H$ of SiNPs suspensions in DMEM supplemented with increasing concentrations of BSA (0.05, 0.1, 0.2, 0.4, 0.6, 0.8, 1, and 2% w/v) were measured by DLS using a Malvern Zetasizer ZS instrument (Malvern Instruments Ltd., UK). Measurements were performed at 25 °C, and reported values represent the average of three independent measurements.

***Cell Viability of RAW 264.7***

Cell viability experiments were performed using RAW 264.7 cells cultured in high-glucose DMEM supplemented with 10% FBS and 1% penicillin/streptomycin. Cells were seeded at a density of 20,000 cells/well in 96-well plates. After 24h, SiNPs at concentrations of 0.3, 0.03, and 0.003 mg/mL were incubated for 8 minutes with 1% bovine serum albumin (BSA) in DMEM. After that, the cells were treated with the nanoparticles for 1 hour at 37 °C, and with 25% dimethyl sulfoxide (DMSO) as a positive control for cell death. Subsequently, these cells, together with the control were washed with PBS. At the end of the treatment, cell viability was assessed at 0 h (post-internalization), 9 h (first doubling time), and 18 h (second doubling time) using the Alamar Blue® colorimetric assay (10% in DMEM). Fluorescence was measured on a Varioskan Lux reader (560/590 nm).

***Flow cytometry***

RAW 264.7 cells were plated at 200,000 cells/well in 6-well plates. After 24h, SiNPs at 0.3, 0.03, and 0.003 mg/mL were pre-incubated for 8 min with 1% BSA in DMEM, then applied to cells for 1 h at 37 °C. After treatment, cells and controls were washed, trypsinized, centrifuged at 300 g for 5 min, washed with 0.01 M PBS, and fixed in cold 70% ethanol for 20 min. This procedure was repeated for 9 h and 18 h post-incubation to assess the effect of cell doubling time (DT). Prior to analysis, cells were washed with PBS, permeabilized, and stained with propidium iodide (PI) solution containing 0.1% Triton X-100, 200 μg/mL RNase A, and 20 μg/mL PI in 0.01 M PBS for 30 min at room temperature. Samples were kept on ice and protected from light until flow cytometry analysis using a BD FACSCanto II. Data were collected using an argon laser (488 nm) for forward and side scatter (FSC, SSC), PI fluorescence (PE detector, 585/42 nm), and ATTO-633 fluorescence (APC detector, 650/60 nm) to confirm nanoparticle internalization.

*Apoptosis*

RAW 264.7 cells were cultured in high-glucose DMEM supplemented with 10% fetal bovine serum (FBS) and 1% penicillin/streptomycin. Cells were then seeded at a density of 200,000 cells per well in 6-well plates. After 24h, SiNPs at a concentration of 0.3 mg/mL were incubated with 1% BSA in DMEM for 8 min. Subsequently, cells were treated with nanoparticles for 1 h at 37 °C. DMSO (25%) was used as a positive control for apoptosis after 24 h of incubation. An Annexin V–Alexa Fluor 488 and propidium iodide (PI) apoptosis detection kit was used to label apoptotic and necrotic cells, following the manufacturer's instructions. Samples were centrifuged, washed with cold 1× PBS, and resuspended in 100 µL of 1× binding buffer. Subsequently, 5 µL of Annexin V and 1 µL of diluted propidium iodide were added, and the samples were incubated for 15 min at room temperature, protected from light. After incubation, 400 µL of binding buffer were added, and samples were kept on ice until analysis. Flow cytometry analyses were performed using a BD FACSCanto II cytometer. Unstained cells were also analyzed as a control condition. This staining procedure was repeated at 9 h and 18 h after nanoparticle internalization.

*Scanning Electron Microscopy (SEM)*

To investigate the nuclear internalization observed, a decellularization protocol was performed based on literature procedures. To validate membrane lysis while minimizing nuclear damage, cells were treated with 1% Triton for different incubation times and imaged by SEM. RAW 264.7 cells were cultured on glass coverslips placed in 6-well plates at a density of 25,000 cells per well. After 24h, samples were treated with 1% Triton for 2, 3, 4, or 6 min, washed twice with PBS, and fixed with 2.5% glutaraldehyde in 0.1

M cacodylate buffer containing 3 mM $CaCl_2$ for 5 min at room temperature, followed by overnight incubation at 4 °C. After fixation, samples were washed three times with 0.1 M cacodylate buffer containing 3 mM $CaCl_2$ for 5 min each. Dehydration was performed using a graded ethanol series (15%, 30%, 50%, 70%, 80%, and 90%, 10 min per step, on ice), followed by a final incubation in 100% ethanol. Samples were dried by $CO_2$ critical point drying, mounted on metal stubs, and analyzed using a scanning electron microscope (Inspect 50, Thermo Fisher Scientific).

#### *Time-Lapse Fluorescence Microscopy*

RAW 264.7 cells were seeded at a density of 100,000 cells per well in 36 × 10 mm dishes containing a glass coverslip at the bottom. After 24h, cells were incubated with fluorescent SiNPs at a concentration of 0.3 mg/mL previously treated with BSA. Time-lapse imaging was then performed using an inverted fluorescence microscope equipped with a spinning disk confocal system at INFABiC (UNICAMP). During image acquisition, cells were maintained under physiological conditions using a microscope-mounted incubation chamber at 37 °C with a controlled atmosphere of 5% $CO_2$ and appropriate humidity. Images were acquired over 1 h with 2 min intervals between frames. After this period, dishes were washed with PBS to remove excess nanoparticles, and cells were returned to the microscope for a second acquisition over 18 h, with images recorded every 15 min in a selected field of view.

**Morphology and Size Distribution**

The synthesis procedure proved to be effective in producing silica nanoparticles with a predominantly spherical morphology, as evidenced by the STEM micrograph shown in Figure S1a. The nanoparticles exhibit well-defined contours and a relatively uniform shape, indicating good control over the nucleation and growth processes during synthesis. No significant population of irregularly shaped or highly aggregated particles was observed, supporting the reproducibility and robustness of the synthetic approach.

The particle size distribution of the SiNPs (Figure S1b) was quantitatively evaluated using ImageJ software through the analysis of STEM images, based on the measurement of 250 individual nanoparticles. From this statistical analysis, an average particle diameter (mean ± standard deviation) of 135.5 ± 16 nm was obtained. The relatively narrow size distribution reflects moderate polydispersity, which is desirable for biological and physicochemical studies. Importantly, most of the nanoparticles exhibited diameters below 200 nm, a size regime that is highly relevant for biological applications, particularly for efficient cellular uptake, intracellular trafficking, and quantitative internalization studies [1].

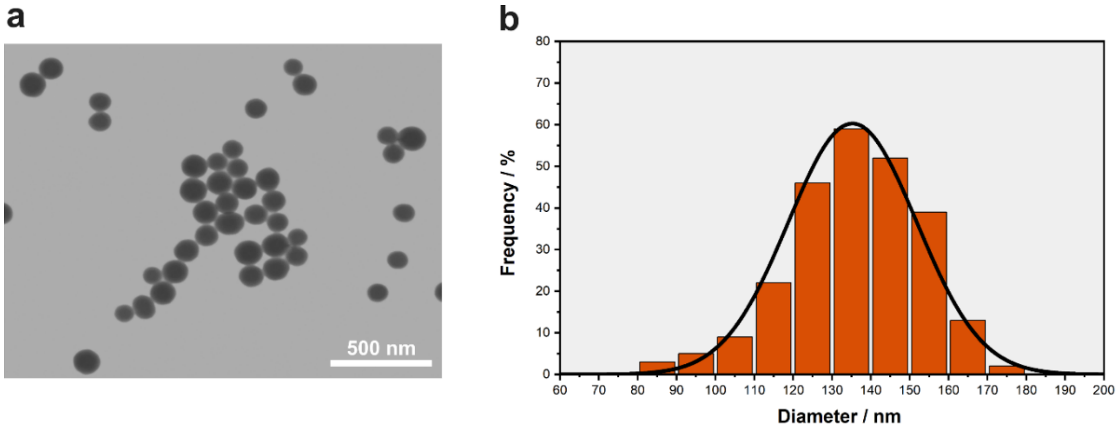


**Figure S1.** a) Micrographs of SiNPs obtained by STEM and b) its size distribution.

**Fluorescent silica nanoparticles**

The selection of ATTO 633 as a fluorescent marker was motivated by its favorable photophysical properties, particularly its absorption and emission at longer wavelengths, which significantly reduce interference from cellular autofluorescence. In addition, ATTO 633 exhibits a high fluorescence quantum yield, as well as excellent thermal and photostability, making it well suited for long-term imaging and quantitative fluorescence analyses.

Following centrifugation of the fluorescent SiNPs suspension, fluorescence emission spectra were recorded separately for the resulting precipitate and the supernatant (Figure S2). The precipitated SiNPs exhibited a strong emission peak centered at approximately 660 nm, characteristic of ATTO 633, whereas the supernatant displayed negligible fluorescence intensity across the measured spectral range. The absence of a detectable signal in the supernatant indicates that no significant dye release occurred, confirming the stable entrapment of ATTO 633 within the silica nanomaterial.

The successful incorporation of the fluorophore into the silica matrix was further supported by the fluorescence spectrum of the ATTO 633–labeled silica nanoparticles shown in Figure S2. To obtain fluorescent SiNPs, a silanized ATTO 633 derivative was introduced into the reaction mixture immediately prior to the addition of TEOS. This synthetic strategy promotes covalent integration of the fluorophore during nanoparticle formation, favoring a homogeneous distribution throughout the nanoparticle volume rather than surface-restricted labeling.

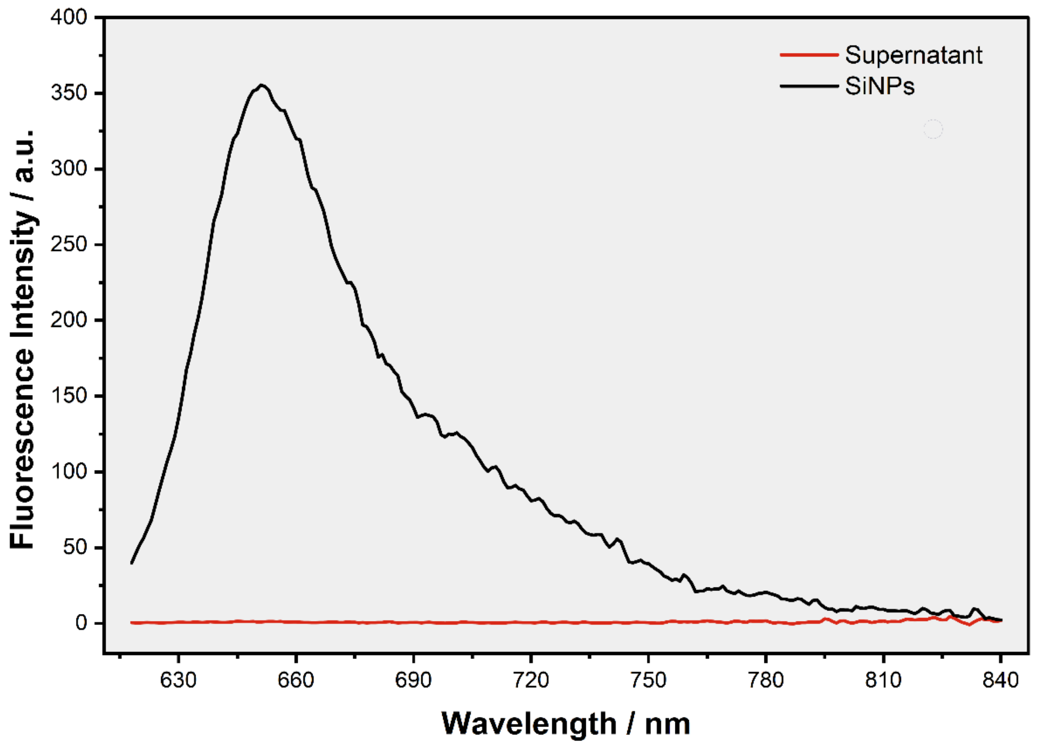


**Figure S2.** The fluorescence spectrum of the precipitate and supernatant obtained after centrifuging a suspension of fluorescent SiNPs.

**Protein Corona Formation**

As shown in Figure S3, the hydrodynamic diameter ($D_H$) of SiNPs increases progressively with increasing BSA concentration in DMEM (0.05–2% w/v), reaching ~520 nm at 2% BSA. This monotonic trend indicates concentration-dependent protein adsorption and progressive corona formation. The absence of abrupt size transitions suggests a continuous adsorption process and a slight aggregation of the nanoparticles.

The comparative data summarized in Table S1 further clarify the role of the biological medium. In water, SiNPs exhibit a $D_H$ of 160.9 nm and a PDI of 0.012, confirming high monodispersity. In protein-free DMEM, the $D_H$ increases dramatically to 1466.6 nm (PDI = 0.302), consistent with strong aggregation driven by ionic screening and reduced electrostatic stabilization. Upon addition of 1% BSA, the $D_H$ decreases to 496.6 nm (PDI = 0.173), demonstrating partial restoration of colloidal stability. Together, these results indicate that while DMEM promotes aggregation, BSA adsorption mitigates this effect through steric stabilization associated with protein corona formation, thereby modulating nanoparticle size under biologically relevant conditions.

.

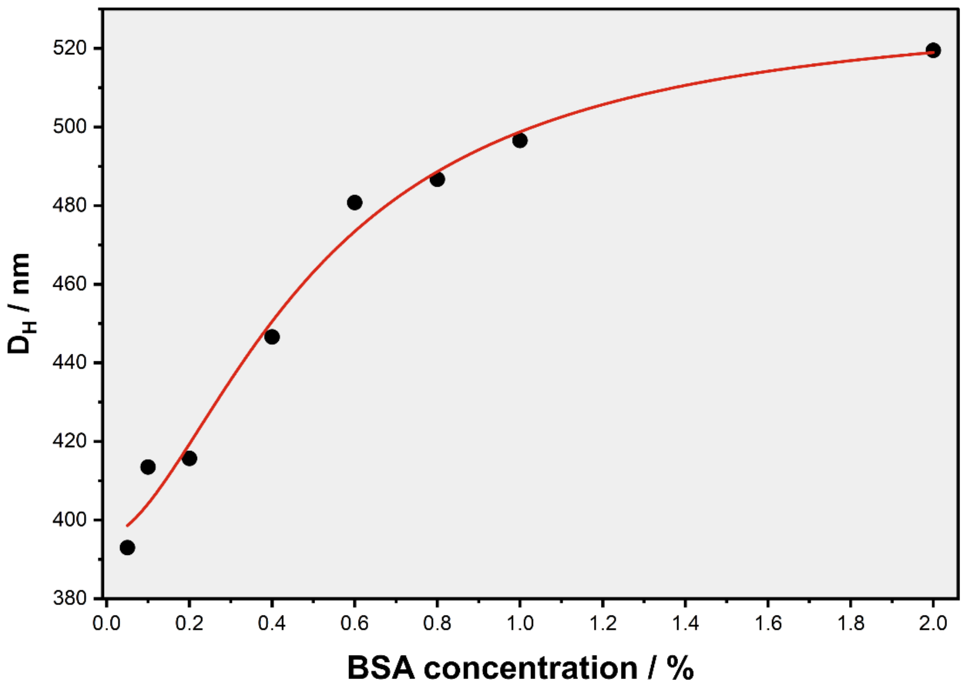


**Figure S3.** Hydrodynamic diameter of SiNPs measured by DLS in DMEM supplemented with increasing concentrations of BSA (0.05–2% w/v). The progressive increase in $D_H$ indicates protein corona formation and a slight aggregation of the nanoparticles.

**Tabel S1.** Average hydrodynamic diameter ($D_H$) and polydispersity (PDI) of SiNPs in water, DMEM medium and DMEM medium with 1% of BSA.

| | $D_H$/ nm | PDI |
|---|---|---|
| SiNPs - H20 | 160.9 | 0.012 |
| SiNPs - DMEM | 1466.6 | 0.302 |
| SiNPs - 1% BSA in DMEM | 496.6 | 0.173 |

**Cytotoxicity Assessment**

Given the observed nuclear internalization of SiNPs at highest concentration (0.3 mg/mL), in RAW 264.7 cells, cell viability assays were performed to determine whether nanoparticle exposure induced acute cytotoxic effects. Cells were treated with SiNPs at concentrations of 0.3, 0.03, and 0.003 mg/mL in the presence of BSA and analyzed at 0 h (immediately after the 1 h incubation period), 9 h, and 18 h post-incubation. Untreated cells served as a negative control, whereas 25% DMSO was used as a positive control for cell death. The results are shown in Figure S4. As expected, positive control induced a pronounced reduction in cell viability at all time points, confirming assay sensitivity. In contrast, SiNP-treated cells maintained high viability across all tested concentrations and time points. Although a modest decrease (≈10–15%) was observed at some intermediate time points, viability values remained above ~85% and showed no concentration-dependent decline. Importantly, even at the highest concentration, cell viability remained comparable to untreated controls throughout the 18 h observation period. These findings indicate that SiNP exposure does not induce detectable acute cytotoxicity under the experimental conditions employed.

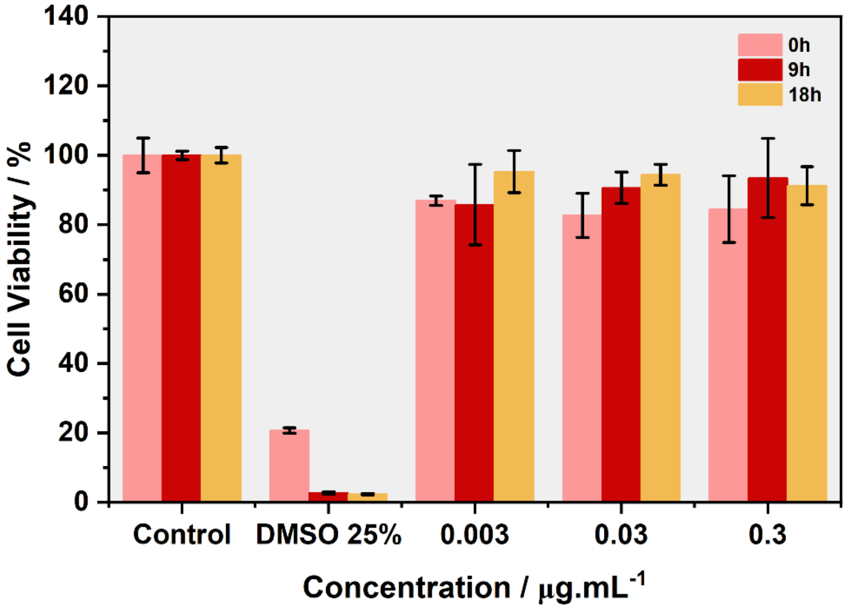


**Figure S4.** Cell viability of RAW 264.7 after incubation with SiNPs at concentrations of 0.3, 0.03, and 0.003 mg/mL in the presence of BSA for 1 h (initial time-0h), 9 h, and 18 h.

**Flow Cytometry Analysis of Nanoparticle Internalization**

To investigate the influence of nanoparticle concentration and cell division on SiNPs distribution in RAW 264.7 macrophages, flow cytometry analysis was performed at three concentrations (0.3, 0.03, and 0.003 mg/mL) and at two subsequent doubling times (9 and 18 h) following a 1 h exposure period (Figure S5). At the initial time point (0 h), high nanoparticle-positive populations were observed for 0.3 and 0.03 mg/mL, with 99% and 98.7% of cells located in the ATTO+PI quadrant, respectively (Figure S5a,b). In contrast, the lowest concentration (0.003 mg/mL) resulted in 79.6% ATTO-positive cells (Figure S5c), indicating concentration-dependent internalization efficiency. After one doubling time (9 h), the percentage of ATTO-positive cells remained high for 0.3 and 0.03 mg/mL (99.3% for both concentrations; Figure S5d,e), demonstrating sustained intracellular retention. However, at 0.003 mg/mL the ATTO-positive population decreased to 64.6% (Figure S5f), suggesting progressive dilution of the nanoparticle load during cell division. Following two doubling times (18 h), cells treated with 0.3 and 0.03 mg/mL still exhibited high levels of nanoparticle positivity (98.9% and 98.5%, respectively; Figure S5g,h), indicating that intracellular nanoparticle levels remain above the detection threshold despite proliferation. In contrast, the lowest concentration (0.003 mg/mL) showed a further reduction to 49.6% ATTO-positive cells (Figure S5i), consistent with mitotic dilution effects. RAW 264.7 macrophages possess multiple uptake pathways, including endocytosis, macropinocytosis, and phagocytosis[2], which likely contribute to the high internalization efficiency observed at the two higher concentrations. Overall, the flow cytometry data demonstrates a clear concentration-dependent persistence of intracellular SiNPs, with nanoparticle retention exceeding two cell doubling cycles at $\geq$0.03 mg/mL whereas lower exposure levels become progressively diluted through cell proliferation.

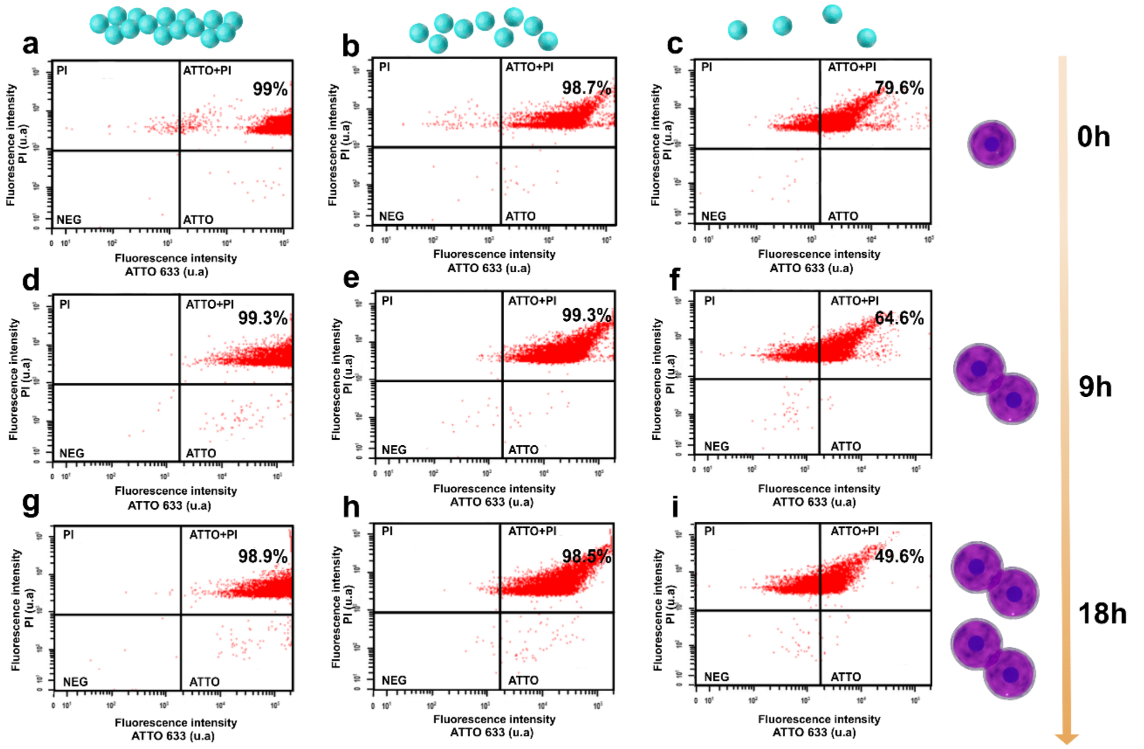


**Figure S5.** Internalization of SiNPs at different concentrations in RAW 264.7 cells, analyzed by flow cytometry. Nanoparticles at the following concentrations and time points: (a) 0.3 mg/mL at 0 h (after 1 h of incubation), (b) 0.03 mg/mL at 0 h, (c) 0.003 mg/mL at 0 h, (d) 0.3 mg/mL at 9 h (first doubling time), (e) 0.03 mg/mL at 9 h, (f) 0.003 mg/mL at 9 h, (g) 0.3 mg/mL at 18 h (second doubling time), (h) 0.03 mg/mL at 18 h, and (i) 0.003 mg/mL at 18 h.

**Apoptosis Analysis by Flow Cytometry**

To further evaluate whether nuclear internalization of SiNPs induces cell death pathways, apoptosis was assessed by flow cytometry using Annexin V/PI staining (Figure S6). Annexin V binds to externalized phosphatidylserine, an early marker of apoptosis, whereas PI stains cells with compromised membrane integrity, indicative of advanced membrane damage. Representative dot plots for RAW 264.7 cells exposed to 0.3 mg/mL SiNPs at 0, 9, and 18 h are shown in Figure S6a, c, and e, respectively, alongside the corresponding untreated controls (Figure S6b, d, and f). Quadrant distribution follows the nomenclature indicated in the plots: NEG, ANEXINA, PI, and ANEXINA+PI.

Across all conditions, many events were localized in the NEG quadrant, indicating viable cells. Only minimal populations were detected in the ANEXINA+PI quadrant, corresponding to late apoptotic/necrotic cells. The ANEXINA and PI quadrants showed similarly low percentages, without marked shifts compared to their respective controls. Quantitative analysis of corrected ANEXINA+PI populations, after subtraction of the corresponding control values at each time point, is summarized in Table S2. The percentage of dead cells remained low, with values of 0.5% at 0 h, 1.5% at 9 h, and 1.6% at 18 h for the 0.3 mg/mL condition. These percentages fall within the expected experimental variability for macrophage cultures and do not indicate biologically significant activation of cell death pathways. Overall, the Annexin V/PI analysis demonstrates that exposure to SiNPs at 0.3 mg/mL, even after two cell doubling cycles, does not promote detectable increases in ANEXINA, PI, or ANEXINA+PI populations relative to controls. These findings corroborate the viability data (Figure S4) and further support that nuclear localization of SiNPs does not induce acute cytotoxic or pro-apoptotic effects under the experimental conditions investigated.

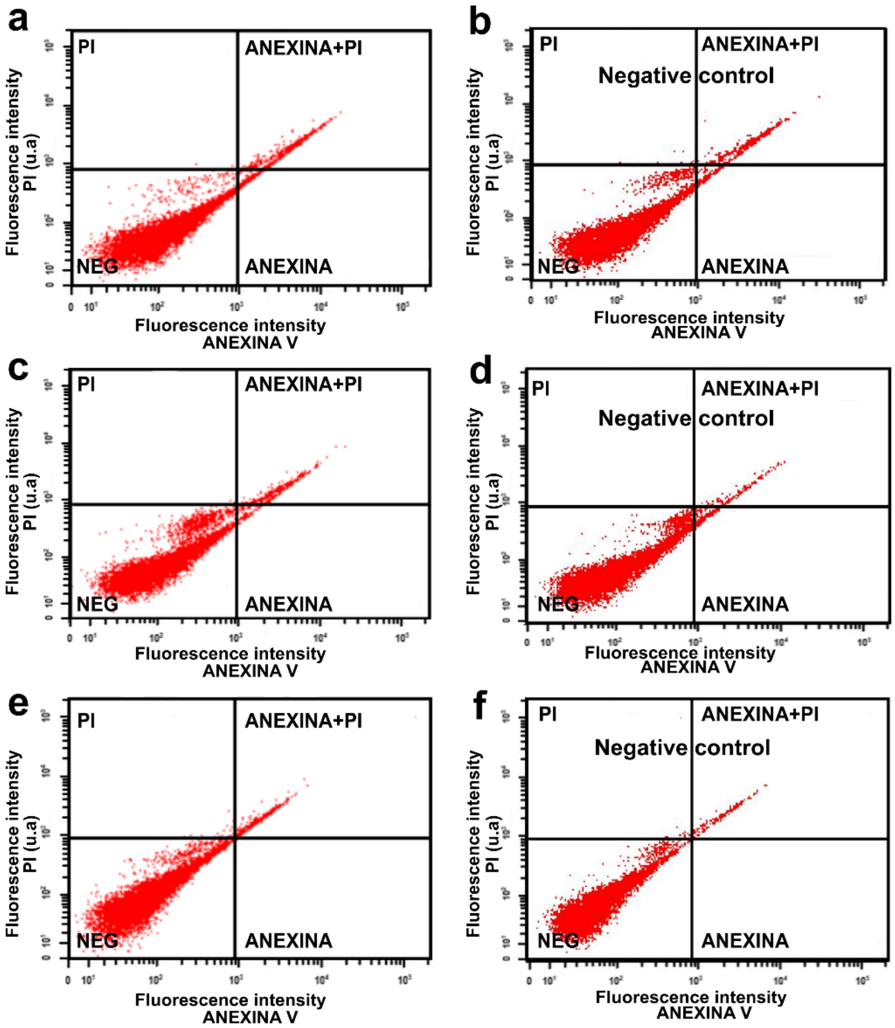


**Figure S6.** Flow cytometry analysis of cell death pathways in RAW 264.7 cells, including early apoptosis (ANEXINA), necrosis (PI), late apoptosis/secondary necrosis (ANEXINA+PI), and viable cells (NEG), under the following conditions: (a) 0.3 mg/mL at 0 h (after 1 h of incubation), (c) 0.3 mg/mL at 9 h (first doubling time), and (e) 0.3 mg/mL at 18 h (second doubling time). Apoptosis assay by flow cytometry in RAW 264.7 cells without nanoparticles (control) at the following time points: (b) 0 h, (d) 9 h, and (f) 18 h.

**Tabel S2.** The percentage of dead cells (Annexin + PI) at nanoparticle concentrations of 0.3 mg/mL at the following time points: 0 h, 9 h, and 18 h. *Percentage values were corrected by subtracting the Annexin + PI percentages observed in the corresponding control at each time point.

| SiNPs@ATTO concentration (mg/mL) | Death cells* (%) (Time 0h) | Death cells* (%) (Time 9h) | Death cells* (%) (Time 18h) |
|---|---|---|---|
| 0,3 | 0,5 | 1,5 | 1,6 |

**Development of a Nuclear Isolation Protocol**

A relevant aspect highlighted in this study was the localization of SiNPs in structures consistent with nuclear envelope invaginations at higher concentrations, a phenomenon not observed under lower concentration conditions, as revealed by cryo-SXT. To further investigate this finding, with a specific focus on nuclear internalization by ptychography, a nuclear isolation protocol was developed to preserve nuclear integrity while selectively lysing the plasma membrane[3].

For this purpose, cells were treated with 1% Triton X-100, followed by glutaraldehyde fixation and critical point drying—conditions required for ptychographic imaging. Figure S7 presents micrographs of cells exposed to different incubation times with 1% Triton X-100, aiming to optimize the selective lysis protocol. An incubation time of 2 min (Figure S7a) was insufficient to promote effective plasma membrane lysis. In contrast, longer incubation times of 4 and 6 min (Figure S7c,d), although effective in membrane disruption, resulted in structural collapse and compromised nuclear integrity. An intermediate incubation time of 3 min (Figure S7b) was therefore established as optimal, enabling selective plasma membrane lysis while preserving nuclear morphology. Under these conditions, cytoskeletal structures resistant to detergent treatment, such as actin filaments and microtubules, remained visible[4].

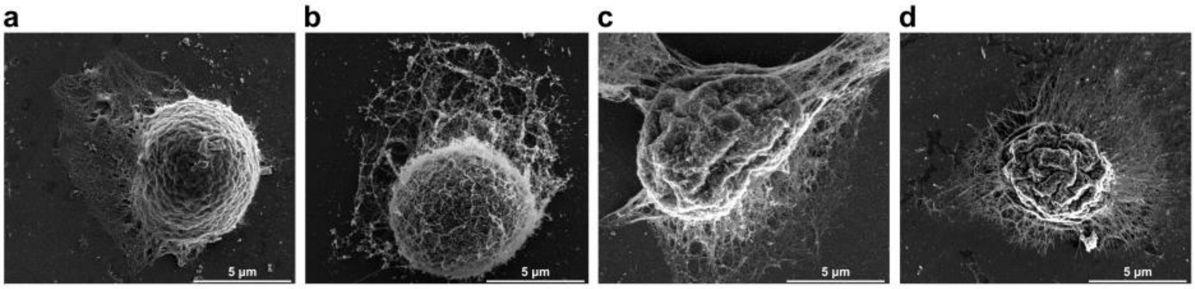


**Figure S7.** SEM micrographs of RAW cells treated with 1% Triton for different incubation times: (a) 2 min, (b) 3 min, (c) 4 min, and (d) 6 min.

### Time-Lapse Fluorescence Microscopy

To investigate the temporal dynamics of nanoparticle mobility, particularly under the highest exposure condition (0.3 mg/mL), time-lapse fluorescence microscopy experiments were performed. Representative frames extracted from the acquired video are shown in Figure S8. During the first hour of incubation (Figure S8a–c), a rapid accumulation of SiNPs (red signal, ATTO633) within RAW 264.7 macrophages was observed under bright-field visualization. Although no specific nuclear staining was performed due to the technical limitations of the inverted spinning disk fluorescence system, a marked increase in intracellular fluorescence intensity was detected throughout the cellular volume, indicating efficient nanoparticle uptake within a short exposure period.

Following nanoparticle incubation, cells were washed with PBS and monitored for an additional 18 h, corresponding to approximately two complete cell division cycles (Figure S8d–f). At early post-wash time points (Figure S8d), the fluorescence signal appeared more diffusely distributed throughout the cytoplasm. Over time, however, a progressive clustering of the fluorescent signal was observed in regions closer to the cellular center (Figure S8e,f). This redistribution pattern suggests active intracellular trafficking of SiNPs into specific subcellular compartments, potentially in the perinuclear region.

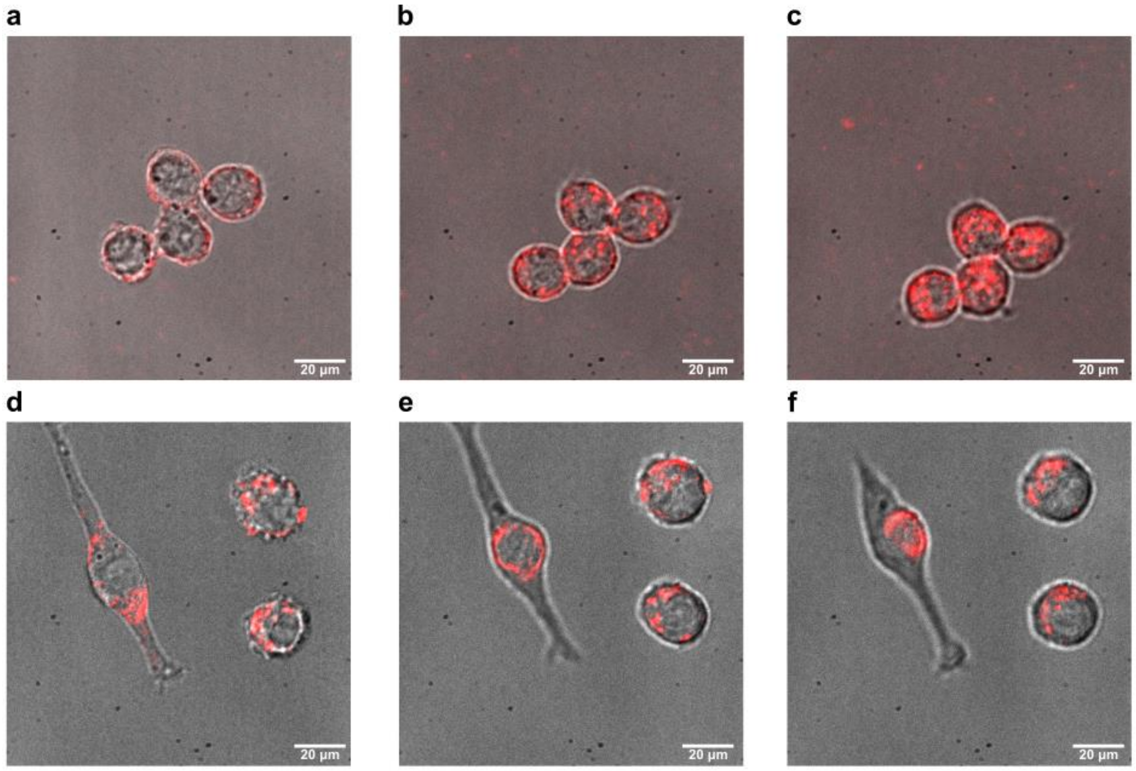


**Figure S8.** Selected frames from time-lapse fluorescence microscopy acquisition (red: SiNPs). Images correspond to the following time points: (a) initial, (b) intermediate, and (c) final during 1 h of incubation with the nanoparticles; and (d) initial, (e) intermediate, and (f) final over 18 h after incubation

### SEM Analysis of Radiation-Induced Structural Alterations

Considering the potential effects of radiation damage on cellular structure, selected silicon nitride membranes used in the experiment were analyzed by SEM, as shown in Figure S9. This complementary imaging approach, performed after the ptychography experiments, enabled important observations that will inform the design of future studies. In Figure S9a, a non-irradiated cell is shown, where the integrity of cytoskeletal filaments, including actin filaments and microtubules, can be clearly identified. In contrast, Figure S9b and S9c, corresponding to cells subjected to the ptychography experiment, exhibit a near-complete absence of these filamentous structures. This indicates that they were degraded by the incident radiation, which is consistent with their ultrastructural characteristics, as they are thin, protein-based components known to be highly radiation-sensitive. These observations support the hypothesis that the structures preserved in the ptychographic images correspond primarily to the isolated cell nucleus and the internalized nanoparticles. Regarding the possibility of radiation-induced nuclear collapse, no visually significant reduction in nuclear volume was observed. However, to increase analytical rigor, the development of a quantitative protocol to compare nuclear diameters before and after radiation exposure is planned for future experiments.

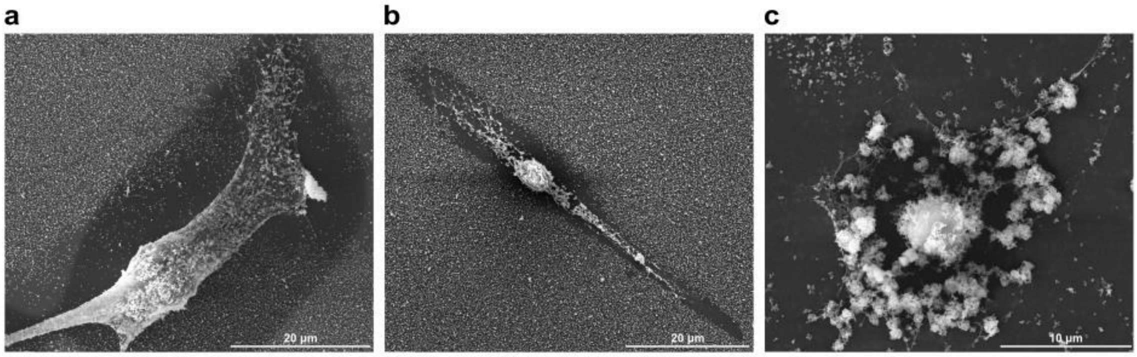


**Figure S9.** SEM micrographs of RAW cells treated with 0.3 mg/mL SiNPs: (a) after 1 h of incubation, without exposure to ptychography radiation; (b) after 1 h of incubation, after ptychography imaging; and (c) after 9 h of incubation, after ptychography imaging with radiation exposure.

**Concentration-Dependent Intracellular Distribution of SiNPs**

Figure S10 presents rendered and segmented tomographic volumes highlighting the intracellular localization of SiNPs in RAW 264.7 cells as a function of nanoparticle concentration. The segmentation enables clear visualization of the spatial relationship between the nucleus (pink), vesicular compartments (gold), and SiNPs (cyan), providing structural context beyond conventional projection images.

At the lowest concentration (0.003 mg/mL, Figure S10a), only a limited number of nanoparticles are detected inside the cell, appearing mostly as isolated structures with sparse association with intracellular vesicles. Increasing the concentration to 0.03 mg/mL (Figure S10b) results in a marked increase in vesicle-associated nanoparticles, indicating active uptake and accumulation within endocytic compartments distributed throughout the cytoplasm. At the highest concentration (0.3 mg/mL, Figure S10c), a pronounced clustering of SiNPs-containing compartments is observed, with several segmented structures localized within the nuclear region. This observation is consistent with the nuclear internalization described in the main text (Figure 2b), from which these segmentations were derived. Rather than representing exclusively perinuclear accumulation, the reconstructed volumes reveal nanoparticle-associated vesicular structures positioned inside the nuclear boundary, supporting the interpretation that a fraction of SiNPs translocates into the nucleus. The increased local nanoparticle density further enhances segmentation contrast, enabling clearer visualization of intracellular compartments in the three-dimensional reconstructions.

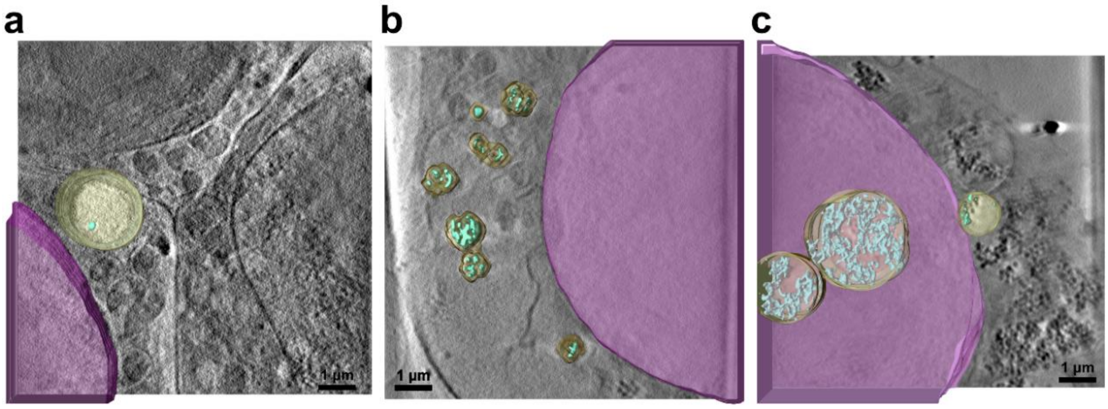


**Figure S10.** Rendered and segmented volumes illustrating the internalization and intracellular distribution of SiNPs in RAW 264.7 cells at different concentrations: a) 0.003 mg/mL, b) 0.03 mg/mL, and c) 0.3 mg/mL. Highlighted structures include the cell nucleus (pink), vesicles (gold), and SiNPs (cyan).

**FSC-Based Resolution Estimation of Ptychographic Tomographic Reconstructions**

The spatial resolution of the three-dimensional tomographic reconstructions obtained by PXCT was quantitatively evaluated using Fourier shell correlation (FSC), as shown in Figure S11. The resolution was determined using the 1/2-bit threshold criterion, which provides a conservative and widely accepted estimate of the reproducible spatial frequency content in 3D reconstructions. The control condition (Figure S11a) exhibited a resolution of approximately 106 nm, consistent with the expected performance of the experimental setup under standard imaging conditions. Upon exposure to SiNPs at 0.3 mg/mL, similar resolutions were obtained after 1 h (Figure S11b) and 9 h (Figure S11c), indicating that nanoparticle internalization alone does not significantly alter reconstruction quality. Notably, the reconstruction corresponding to 18 h incubation (Figure S11d) displayed an improvement in resolution, reaching ~84 nm. This enhancement can be attributed to the increased intracellular aggregation of nanoparticles observed at later time points. In ptychographic imaging, reconstruction fidelity strongly depends on the strength and diversity of the recorded diffraction signal. Aggregated nanoparticles act as high electron-density scattering centers, increasing phase contrast and improving the signal-to-noise ratio at higher spatial frequencies. The presence of stronger scattering features introduces additional constraints during iterative phase retrieval, thereby stabilizing convergence and enabling recovery of finer structural details.

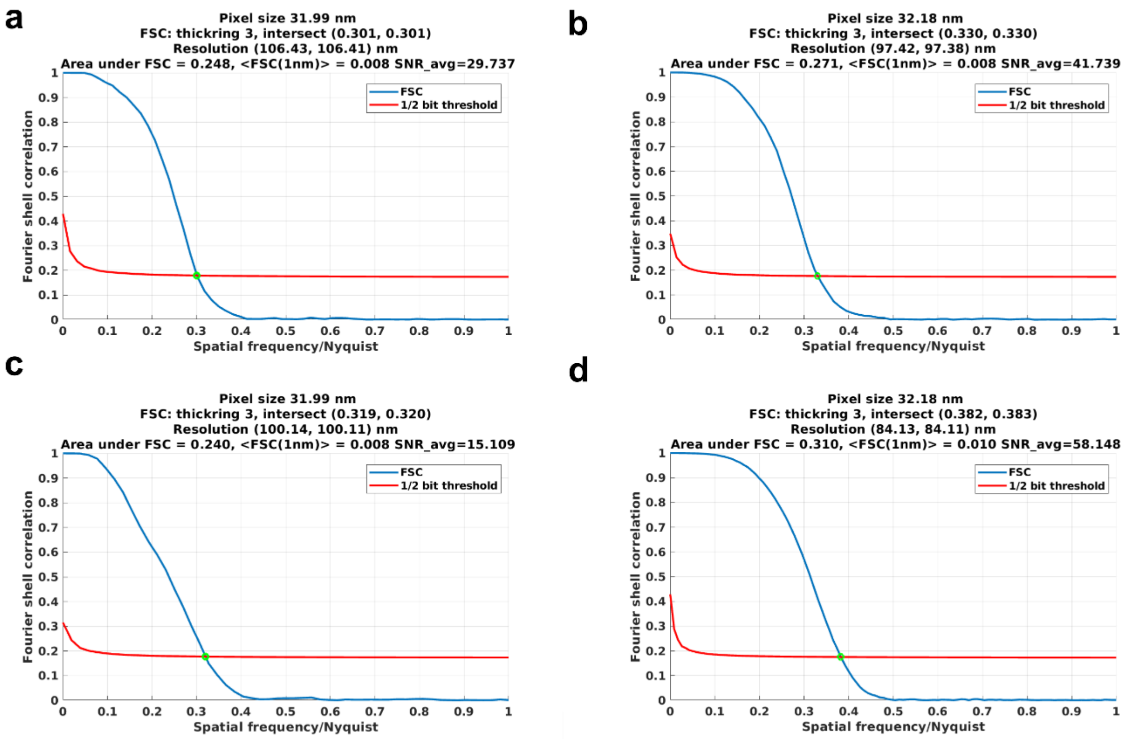


**Figure S11.** Fourier shell correlation (FSC) analysis used to estimate the spatial resolution of the three-dimensional ptychographic tomographic reconstructions obtained under different experimental conditions. (a) Control condition corresponding to the reconstruction shown in Figure 6d (main text). (b) Cells exposed to SiNPs at 0.3 mg/mL after 1 h of incubation (Figure 6h). (c) Cells exposed to 0.3 mg/mL after 9 h (first doubling time; Figure 6l). (d) Cells exposed to 0.3 mg/mL after 18 h (second doubling time; Figure 6p). Resolution values were determined using the 1/2-bit FSC criterion.

**Legends of supplementary videos**

**Movie_S1.** Tomographic reconstruction and segmentation of a RAW 264.7 cell obtained by cryo-SXT after 1 h incubation with SiNPs (0.3 mg/mL).

**Movie_S2**. Tomographic reconstruction and segmentation of RAW 264.7 cell obtained by cryo-SXT exposed to SiNPs (0.03 mg/mL) at the initial time point

**Movie_S3** Tomographic reconstruction and segmentation segmentation of RAW 264.7 RAW 264.7 cell obtained by cryo-SXT exposed to SiNPs (0.003 mg/mL) at the initial time point.

**Movie_S4**. Tomographic reconstruction and segmentation of RAW 264.7 cell obtained by cryo-SXT exposed to SiNPs (0.3 mg/mL) at the first doubling time.

**Movie_S5**. Tomographic reconstruction and segmentation of RAW 264.7 cell obtained by cryo-SXT to SiNPs (0.3 mg/mL) at the second doubling time.

**Movie_S6.** Ptychographic 3D segmentation of a RAW 264.7 macrophage exposed to SiNPs (0.3 mg/mL) at the initial time point.

**Movie_S7**. Ptychographic 3D segmentation of a RAW 264.7 macrophage exposed to SiNPs (0.3 mg/mL) at the first doubling time.

**Movie_S8.** Ptychographic 3D segmentation of a RAW 264.7 macrophage exposed to SiNPs (0.3 mg/mL) at the second doubling time.
